# In situ structures of periplasmic flagella reveal a distinct cytoplasmic ATPase complex in *Borrelia burgdorferi*

**DOI:** 10.1101/303222

**Authors:** Zhuan Qin, Akarsh Manne, Jiagang Tu, Zhou Yu, Kathryn Lees, Aaron Yerke, Tao Lin, Chunhao Li, Steven J. Norris, Md A. Motaleb, Jun Liu

**Affiliations:** Department of Microbial Pathogenesis & Microbial Sciences Institute, Yale University, New Haven, CT 06519; Department of Pathology and Laboratory Medicine, McGovern Medical School, Houston, TX 77030.; Department of Microbiology and Immunology, Brody School of Medicine, East Carolina University, Greenville, NC 27834; Philips Research Institute, School of Dental Medicine, Virginia Commonwealth University, Richmond, VA 23298

## Abstract

Periplasmic flagella are essential for the distinct morphology and motility of spirochetes. A flagella-specific Type III secretion system (fT3SS) composed of a membrane-bound export apparatus and a cytosolic ATPase complex is responsible for the assembly of the periplasmic flagella. Here, we combine cryo-electron tomography and mutagenesis approaches to characterize the fT3SS machine in the Lyme disease spirochete *Borrelia burgdorferi*. We define the fT3SS machine by systematically characterizing mutants lacking key component genes. We discover that a distinct cytosolic ATPase complex is attached to the flagellar C-ring through multiple spoke-like linkers. The ATPase complex not only strengthens structural rigidity of the C-ring, but also undergoes conformational changes in concert with flagellar rotation. Our studies provide structural framework to uncover the unique mechanisms underlying assembly and rotation of the periplasmic flagella and may provide the bases for the development of novel therapeutic strategies against several pathogenic spirochetes.

## INTRODUCTION

Spirochetes are a group of bacteria responsible for several serious human diseases including Lyme disease (*Borrelia* or *Borreliella* species), syphilis (*Treponema pallidum* subsp. *pallidum*), and leptospirosis *(Leptospira interrogans* and other *Leptospira* species). Spirochetes are easily recognized by their distinctive wave-like or helical morphology and unique form of motility, but remain poorly understood. Their motility is driven by periplasmic flagella, which reside and rotate within the periplasmic space between the outer membrane and the peptidoglycan layer. Recent genetic studies indicate that their unique motility is crucial for host infection and/or bacterial transmission by all pathogenic spirochetes studied to-date (Lambert et al., 2012; Li et al., 2010; Motaleb et al., 2015; Sultan et al., 2013; Sultan et al., 2015; Wunder et al., 2016). Rotation of the periplasmic flagella against the elastic cell body contributes to the distinctive morphology and motility of spirochetes. For example, mutants *B. burgdorferi* that lack their periplasmic flagellar filaments encode by FlaB are non-motile and rod shaped (Charon et al., 2012; Charon et al., 2009; Motaleb et al., 2000; Sultan et al., 2013; Sultan et al., 2015).

The periplasmic flagella are different from other bacterial flagella in many aspects. In particular, the spirochetal flagellar motor is significantly larger than the motors from most other bacteria, including those of the model organisms *Escherichia coli* and *Salmonella enterica* (~80 *vs* ~45 nm in diameter). Unique spirochete-associated features include the periplasmic collar, which is prominent in *B. burgdorferi* (Moon et al., 2016) and all other spirochetes characterized to date (Chen et al., 2011; Liu et al., 2010a; Murphy et al., 2006; Raddi et al., 2012). The large spirochetal flagellar motors produce the highest torque (~4000 pN.nm) observed in bacteria (Beeby et al., 2016). In addition, spirochetes have unusual flagellar hooks in which the hook proteins are cross-linked by a covalent bond that is required to transmit the torque from the motor to the filament (Miller et al., 2016). These spirochete-specific features enable the spirochetes to bore through viscous environments in the hosts.

The filament is the largest component of the periplasmic flagella; it is often as long as 10 μm. Multiple filaments arising from both poles form flat ribbons that wrap around the cell body in a righthanded fashion (Charon et al., 2009). The assembly of the periplasmic flagella is a finely orchestrated process, which includes the initiation of the motor complex subterminally at the cell poles and the formation of the hook and the filament in the periplasmic space (Zhao et al., 2013). The flagellar-specific type III secretion system (fT3SS) is responsible for the assembly of the periplasmic flagella. Although periplasmic flagella are different in location from the external flagella such as those seen in *E. coli*, the fT3SS is conserved among different bacterial species (Chen et al., 2011; Zhao et al., 2014). Furthermore, the fT3SS is evolutionally related to the virulence (v)T3SSs that promote bacterial virulence by delivering effector proteins into eukaryotic cells (Diepold and Armitage, 2015; Erhardt et al., 2010). The secretion process of T3SS is energized by proton motive force (PMF) (Erhardt et al., 2014; Minamino and Namba, 2008; Paul et al., 2008) and ATP hydrolysis (Claret et al., 2003; Fan and Macnab, 1996; Imada et al., 2007). However, the mechanisms underlying the secretion process are not well understood at molecular level.

The fT3SS consists of a membrane-bound export apparatus and a large cytosolic ATPase complex. Six transmembrane proteins (FlhA, FlhB, FliO, FliP, FliQ, and FliR) are thought to form the export apparatus for substrate secretion. Among the six integral membrane proteins, FlhA is the largest, consisting of an N-terminal domain with eight transmembrane regions (FlhA_TM_) and a C-terminal cytoplasmic domain (FlhA_C_) (Macnab, 2003; McMurry et al., 2004). FlhA facilitates the translocation of the export substrates into the central channel of the growing flagella (Kihara et al., 2001; Minamino et al., 2010; Zhu et al., 2002). FlhB consists of an N-terminal transmembrane domain (FlhB_TM_) and a C-terminal cytoplasmic domain (FlhB_C_) (Ferris et al., 2005). FliO, FliP, FliQ and FliR are integral membrane proteins, that are also important for substrate secretion (Erhardt et al., 2010).

Three cytoplasmic proteins (FliH, FliI, and FliJ) form the ATPase complex that promotes the export process by binding and delivering substrates to the export apparatus (Fraser et al., 2003; Minamino and Imada, 2015). FliI is an ATPase and shows structural similarity with the α and β subunits of the F_0_F_1_-ATP synthase (Ibuki et al., 2011). FliI exhibits its full ATPase activity when it self-assembles into a homo-hexamer (Imada et al., 2007; Macnab, 2003). FliH probably acts as a negative regulator of the FliI ATPase, and FliJ has a chaperone-like activity that prevents substrate aggregation (Fraser et al., 2003). FliH, FliI, and FliJ coordinately deliver a chaperone-substrate complex to the export gate by binding to the docking platform of the fT3SS for substrate export (Abrusci et al., 2013). FliH_2_ binds to FliI ATPase and localizes FliI to the bottom of flagellar motor through the interaction with FliN on the C-ring (Minamino et al., 2009). FlhA is required for stable anchoring of the FliI_6_ ring to the gate (Bai et al., 2014).

Recent cryo-electron tomography (cryo-ET) studies have revealed the overall structures of the fT3SS machine within intact flagellar motors (Abrusci et al., 2013; Chen et al., 2011; Liu et al., 2010a; Liu et al., 2009; McMurry et al., 2006; Raddi et al., 2012; Zhao et al., 2013). However, these studies have yet provided sufficient details to dissect protein-protein interactions at the molecular level. More importantly, the structure and function of each component have not been systematically analyzed in the context of the intact flagellar motor.

*B. burgdorferi* is the best-studied spirochete system. Recent breakthroughs in genetic manipulations allow the production of well-defined mutations without imposing any secondary alterations (Moon et al., 2016; Motaleb et al., 2011; Sultan et al., 2015; Zhao et al., 2013). The small cell diameter and the highly ordered array of multiple flagellar motors in *B. burgdorferi* make it an excellent model system for *in situ* structural analysis of the periplasmic flagella by cryo-ET. Our previous structural analysis of wild-type cells and several rod mutants of *B. burgdorferi* not only revealed the sequential assembly of the flagellar rod, hook, and filament, but also showed the intact fT3SS machine within the C-ring and beneath the MS-ring (Zhao et al., 2013). Furthermore, disruption of the *fliH* and *fliI* genes by transposon mutagenesis was found to disrupt the assembly and placement of the cytoplasmic ATPase complex and to greatly inhibit flagellar filament formation, which were largely restored by genetic complementation (Lin et al., 2015). However, structural details of the fT3SS machine and its interactions with other flagellar components remain elusive, likely because of the dynamic nature of the fT3SS machine and the difficulty of symmetry-matching among the flagellar subunits.

In this study, we used cryo-ET and sub-tomogram averaging to reveal novel features of the intact fT3SS machine in wild-type (WT) *B. burgdorferi* flagellar motor. We gained an in-depth understanding of the subunit organization and function of the fT3SS machine by systemically characterizing structural changes in several single and multiple deletion variants of the fT3SS in the *B. burgdorferi* flagellar motor. Comparison of these results with recent studies of the T3SSs in external flagella and evolutionarily related bacterial injectisomes provides new insights into these nanomachines that share a common evolutionary origin, but are structurally and functionally different (Hu et al., 2017; Hu et al., 2015; Kawamoto et al., 2013; Zhu et al., 2017).

## RESULTS

### *In situ* analysis of the *B. burgdorferi* flagellar motor reveals novel features of the fT3SS machine

Nine conserved proteins of *B. burgdorferi* are believed to form the membrane-bound export apparatus and the cytosolic ATPase complex (Fig. 1), although the exact details remain to be defined. To dissect the molecular architecture of the intact fT3SS machine, we utilized high-throughput cryo-ET and a sophisticated sub-tomogram classification to analyze the *B. burgdorferi* flagellar motor. By analyzing images of over 20,000 motors extracted from the cell poles, we generated an asymmetric reconstruction that not only reveals the previously observed 16-fold symmetry of the collar and stator structures (Liu et al., 2009; Moon et al., 2016; Zhao et al., 2013) but also discloses a spoke-like structure underneath the C- and MS-rings (Fig. 2A-F, Movie S1). The spoke-like densities extend from a hexagonal “hub” to the bottom of the C-ring (Fig. 2A, B, E). Multivariate statistical analysis (Winkler, 2007) indicates that there are 23 linkers in most flagellar motors, albeit this number can be varied from 21 to 24 in some rare instances (see Fig. S1).

**Figure 1.**
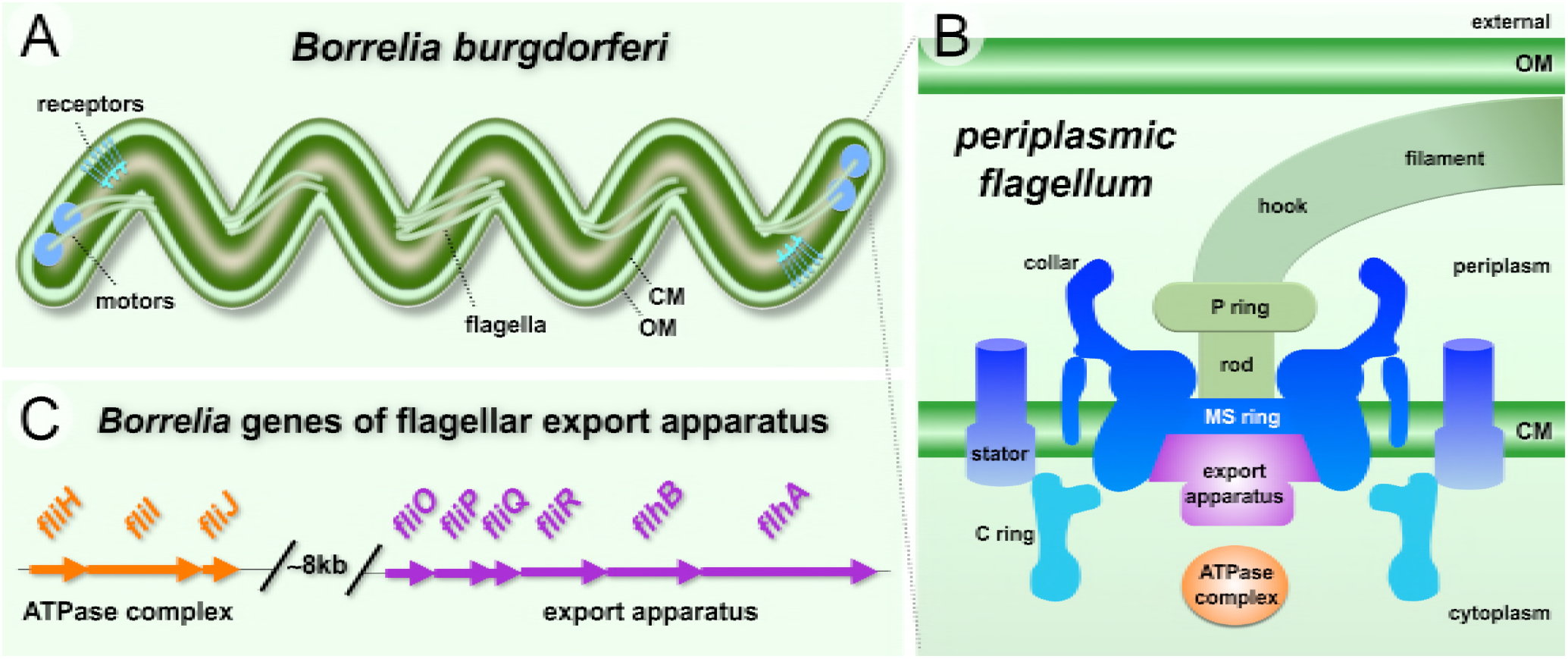
Overview of the fT3SS in *B. burgdorferi*. (A) Schematic model of *B. burgdorferi* and its periplasmic flagella. (B) Model of the spirochete flagellar motor, showing the locations of major components. The fT3SS is localized in the central region of the MS ring and the C-ring. (C) Gene clusters encoding the fT3SS of *B. burgdorferi*. The six genes that encode membrane proteins are marked in purple, while the genes that encode the cytoplasmic proteins are marked in orange. Note that *fliJ* and *fliO* were also called *flbA* and *FliZ*, respectively.

**Figure 2.**
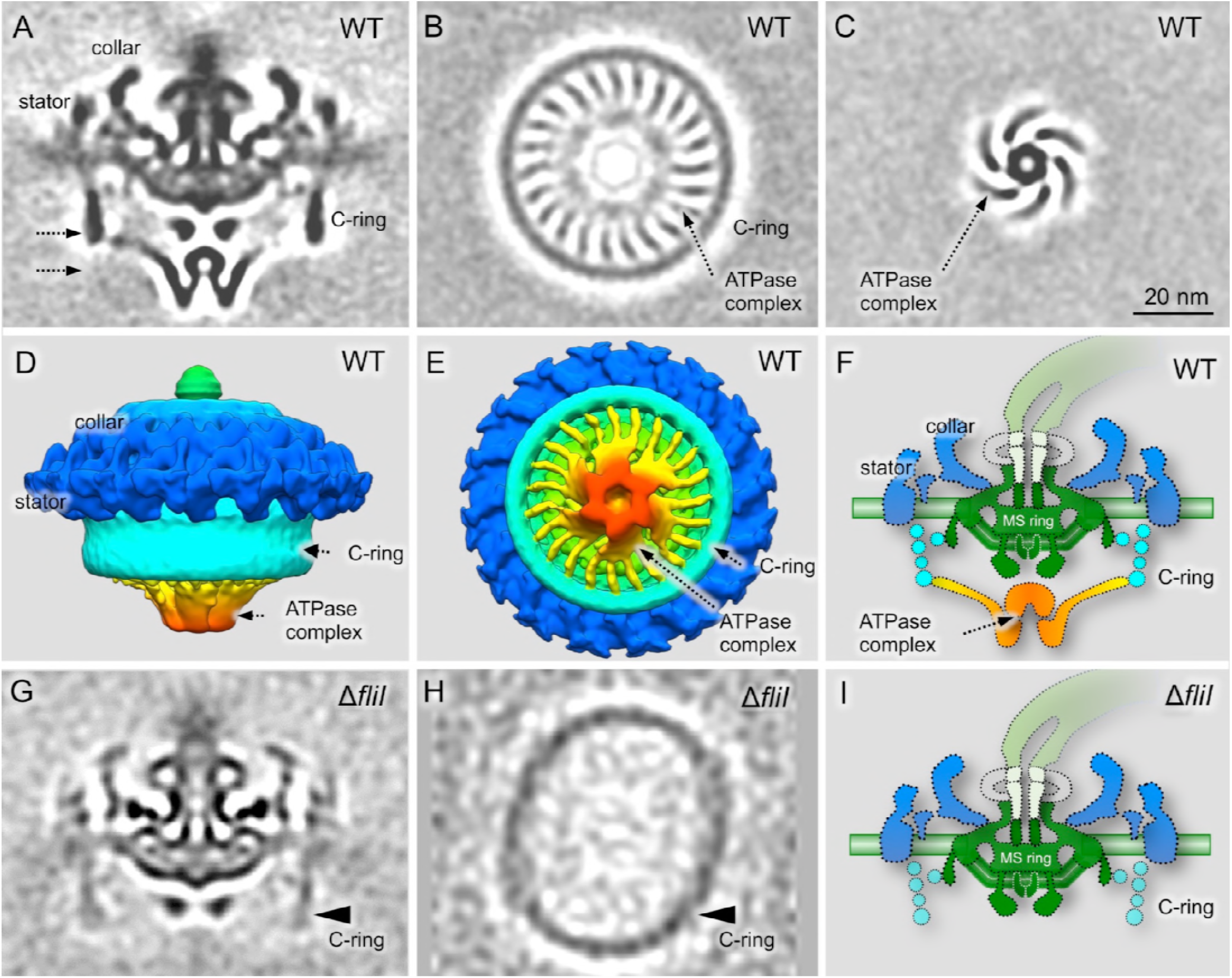
Cryo-ET reveals a novel ATPase complex structure in *B. burgdorferi*. (A) A central section of a flagellar motor structure from WT cells. The structure was generated by alignment of the ATPase ring complex region and classification of the linker region (see S1). (B) A cross section of the C-ring and linker region. 23 linkers connect the ATPase complex to the C-ring. (C) Another cross section shows the hexagonal “hub” structures. (D, E) Surface rendering of the WT flagellar motor structure from a side view and a bottom view, respectively. (F) A schematic model of the WT motor based on the averaged structure showed in (A). (G) A central section of the flagellar motor structure from a *fliI* mutant. The C-ring density from the *fliI* mutant is weak compared to that of the WT. (H) A cross section of the C-ring from the *fliI* mutant shows an ellipse-like structure, which is very different from the C-ring in the WT flagellar motor (B). (I) A schematic model of the flagellar motor structure in the *fliI* mutant.

### The ATPase complex is directly connected with the C-ring

To characterize the ‘hub and spoke’ structures, we compared the structure of the WT motor with those derived from *fliH* and *fliI* mutants (Lin et al., 2015). In either mutant, the ‘hub and spoke’ densities are absent, as previously reported, suggesting that they are formed by the FliI/FliH ATPase complex (Lin et al., 2015). The C-ring density in the average structures from those two mutants is indistinct compared to that derived from the WT motor (Fig. 2G). However, classification shows that the C-ring from the WT maintains a round shape, while the C-ring from the *fliH* mutant is often ellipse-shaped (Fig. 2H). Therefore, we propose that the FliI/FliH complex forms a large ‘hub and spoke’ structure that interconnects with the C-ring and plays an unexpected role in stabilizing this prominent C-ring structure.

### FlhA, FlhB, FliP, FliQ, and FliR are essential for flagellar assembly in *B. burgdorferi*

The large density between the ATPase complex and the MS ring is thought to be the export apparatus. To characterize this density (see Fig. 2A and F), we constructed single Δ*flhA*, Δ*flhB*, Δ*fliP*, Δ*fliQ*, Δ*fliR* mutants, respectively. All five mutants are rod-shaped and non-motile, and they do not form flagellar hook or filament, indicating that FlhA, FlhB, FliP, FliQ, FliR are all essential for flagellar assembly and motility in *B. burgdorferi* (see Fig. S3 for an example). However, the flagellar motors are readily visible at the cell tips of these mutants, largely because of the presence of the periplasmic collar (Moon et al., 2016). The collar is also particularly useful as a reference during the subsequent sub-tomogram averaging. Indeed, sub-tomogram averaging of the Δ*flhA* motors reveals the common feature of the collar, but it also shows that the large, complex density underneath the MS ring is absent (Fig. 3C). Moreover, the cytoplasmic membrane beneath the MS-ring is concave (Fig. 3C), suggesting that reorganization of the membrane components occurred in the absence of FlhA. Moreover, the large donut-shaped density beneath the MS ring is absent in Δ*flhA* (Fig. 3C) comparing with WT (indicated by an arrow in Fig 3F). We thus processed sub-tomogram alignment focusing on the density with WT motors. It turned out that the large donut-shaped density has 9-fold symmetry, and has multiple slim links connecting to MS ring (Fig. 3G, Fig. S2). We speculate the donut shaped density should be FlhAc, as its homolog MxiA has 9-fold symmetry and locates at similar position in injectisome (Abrusci et al., 2013). Similar motor structures were also found in the point mutant *B. burgdorferi flhAD158E* cells that exhibit non-motile phenotype, or the reduced-motility mutant displayed by the *flhA*D158N cells (not shown).

**Figure 3.**
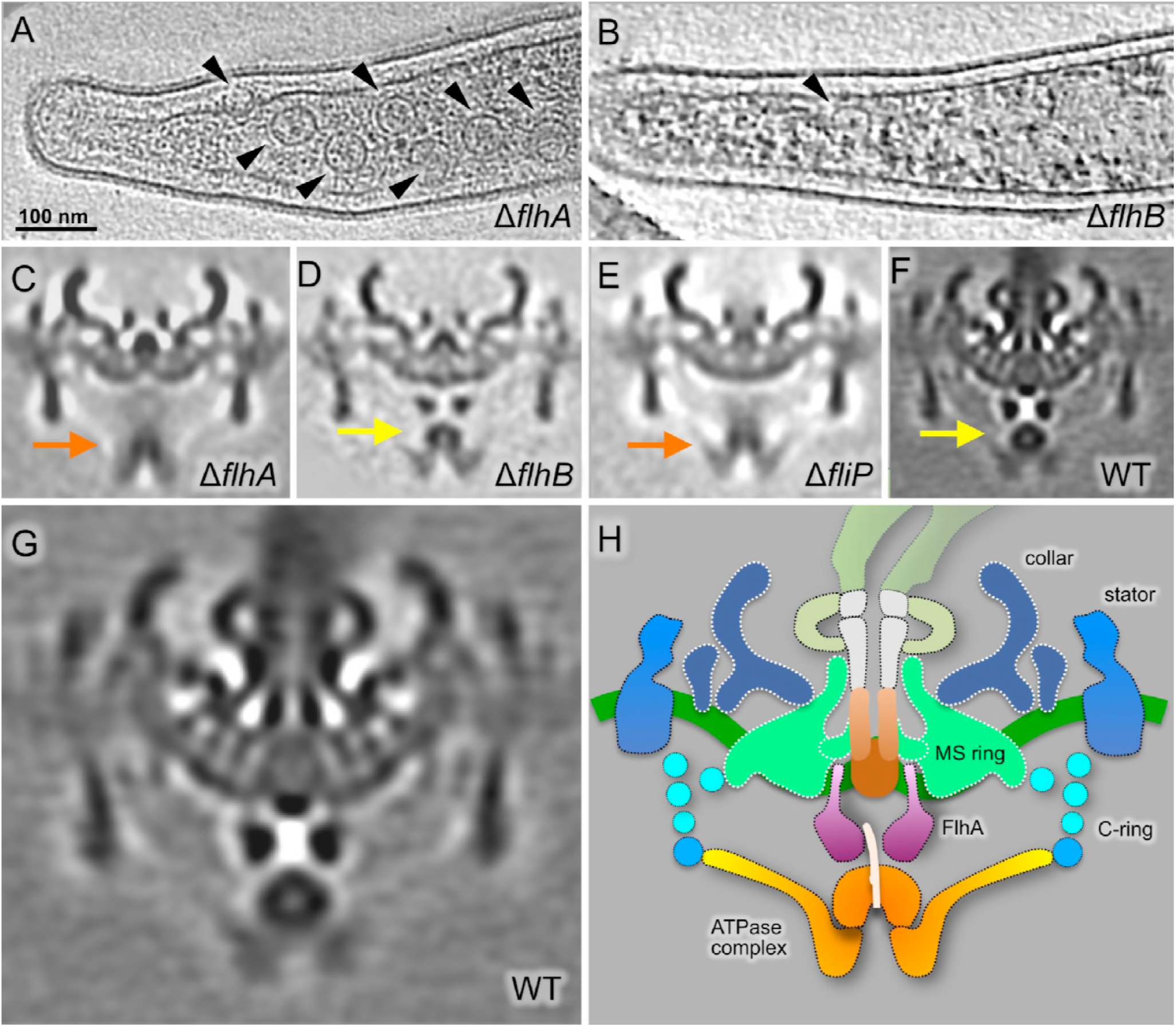
Characterization of the export apparatus. (A) Section of a tomogram from the Δ*flhA* mutant. Several motors can be seen in this representative cell tip (arrowheads). (B) Section of a tomogram from the Δ*flhB* mutant. Only one motor can be seen within this representative cell tip (arrowhead). Central sections of the average density maps of the flagellar motor are shown for (C) the Δ*flhA* mutant; (D) the Δ*flhB* mutant; (E) the Δ*fliP* mutant; and (F) WT. (G) Enlarged view of the average map of the flagellar motor from the WT. (H) Schematic model of the WT *B. burgdorferi* motor showing the proposed locations of fT3SS components in orange-yellow-purple.

In the Δ*flhB* mutant, both the ATPase complex and the export apparatus are evident (Fig. 3D), suggesting that FlhB does not contribute to the structure or positioning of the ATPase complex or the export apparatus. In contrast, in the Δ*fliP* (Fig. 3E, Fig. S4), Δ*fliQ* and *ΔfliR* mutants (Fig. S5, S6), the membrane underneath the MS-ring has a flat surface, and the FlhA complex appears to be absent or disordered. Therefore, the structures of these mutants are strikingly different from the WT structure (Fig. 3F).

Comparative analysis of the four structures shown in Fig. 3C-F suggests that the cytoplasmic domains of the FlhA complexes form the large torus density and the membrane portions of the FlhA complexes are inserted into the cytoplasmic membrane (as proposed in Fig. 3H). It has been recently suggested FliP/Q/R are likely to form the central channel complex (Kuhlen et al., 2018; Ward et al., 2018). Our study provided evidence that the assembly of the FlhA complex depends on the formation of the FliP/Q/R channel. FlhB is not well-defined in our structures, but it is essential for fT3SS function.

### The FlhA complex stabilizes the ATPase complex

The FliI / FliH ATPase complex appears to be associated with the bottom portion of the C-ring, even in the absence of FlhA. However, the FliI/FliH-associated density in the Δ*flhA* mutant is indistinct, indicating that the FlhA complex is involved in stabilization of the ATPase complex under the C-ring (Fig. 3C). In addition, the ATPase complex appears to shift away from the MS-ring, implying that there is an interaction between the FlhA and ATPase complexes. Therefore, we propose that the FlhA complex is not essential for the assembly of the ATPase complex, but it provides a docking site to stabilize the ATPase complex. Our result is consistent with previous study that FlhA is required for stable anchoring the FliI_6_ ring to the export gate (Bai et al., 2014).

### FliO has a limited role in the flagellar assembly in *B. burgdorferi*

FliO is the less conserved among the membrane proteins of the export apparatus. In fact, *B. burgdorferi* FliO and its *Salmonella* homolog have very weak sequence identify (13%; Table S1). We generated a Δ*fliO* mutant, which is less motile than the WT cells. However, cryo-ET reconstructions revealed that both the flagellar motor and filaments are present in the Δ*fliO* mutant (Fig. 4A, F). Furthermore, the flagellar motor and the fT3SS machine in the Δ*fliO* mutant (Fig. 4C) are similar to those in WT, suggesting that FliO is relatively less impo rtant for the formation of the fT3SS and the assembly of the flagellar rod, hook and filament in *B. burgdorferi*.

**Figure 4.**
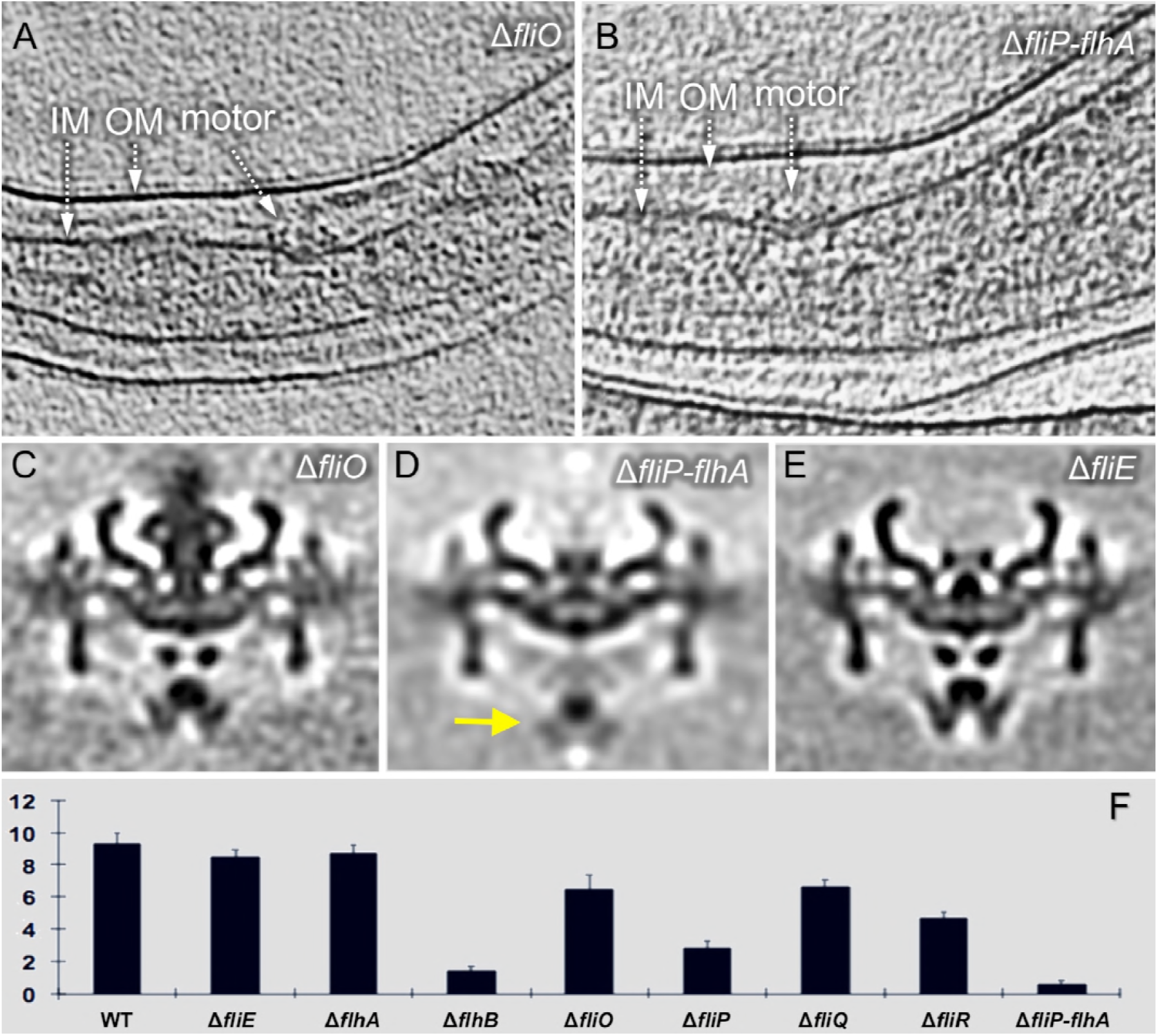
Structural characterization of the fT3SS in a Δ*fliO* mutant and a quintuple mutant. (A) Section of a tomogram of a Δ*fliO* mutant. (B) Section of a tomogram from a quintuple mutant in which the contiguous genes *fliP, fliQ, fliR, flhB*, and *flhA* were deleted. Central sections of average density maps are shown for the flagellar motors from (C) the Δ*FliO* mutant; (D) the quintuple mutant Δ*fliP-flhA*; and (E) the Δ*fliE* mutant, in which the export apparatus is intact while the rod is absent. (F) The number of motors per cell tip in the WT cells and the mutant cells is indicated.

### The export apparatus has a profound impact on flagellar motor formation in *B. burgdorferi*

To understand the overall contribution of the membrane-bound export apparatus proteins to the structure and assembly of the flagellar motor, we generated a quintuple Δ*fliP-flhA* mutant by deleting *fliP, fliQ, fliR, flhB* and *flhA* genes using the Cre-LoxP method (Bestor et al., 2010). As expected, the Δ*fliP-flhA* mutant is also rod-shaped and non-motile. Cryo-ET reconstruction of the mutant shows few motors on the cell tip (Fig. 4B). The average structure of the quintuple-mutant motor (Fig. 4D) is similar to that of the Δ*fliQ* motor (Fig. 4C), although the resolution of the image is relatively poor because fewer motors were available for sub-tomogram averaging. Importantly, the ATPase complex (indicated by orange arrow in Fig. 4D) remains in a similar location as in the Δ*fliQ* or Δ*flhA* motors, supporting the notion that the export apparatus is dispensable for the formation of the ATPase complex. However, absence of the export apparatus proteins has a significant impact on motor formation, as the number of motors per cell tip is highly variable in the mutants of the export apparatus (Fig. 4F). The number of the flagellar motors at each cell tip of the Δ*flhA* mutant is comparable to that of WT (Fig. 4F), suggesting that FlhA does not have a significant effect on motor formation in *B. burgdorferi*. In contrast, the number of the motors in the Δ*flhB* mutant is significantly lower than in WT cells and the Δ*flhA* mutant. These results indicate that FlhB is important for the formation of the motor in *B. burgdorferi* (Fig. 4F), although some motors can still be assembled in the absence of FlhB, FliO, FliP, FliQ, or FliR (Fig. 4F). The impact of the export apparatus proteins on motor formation seems to be cumulative, as demonstrated by the significant decrease in the number of the flagellar motors detected in the quintuple Δ*fliP-flhA* mutant (Fig. 4F). Out of 342 cryo-ET reconstructions from the quintuple mutant, we only identified 54 motors, indicating that the flagellar motors assemble at a very low frequency in the absence of the major membrane proteins (Fig. 4F). Our results are consistent with a model in which there is substantial coordination between the assembly of the MS ring and the export apparatus during the initiation of flagellar assembly (Bai et al., 2014).

### Molecular architecture of the fT3SS machine in *B. burgdorferi*

To better understand the interactions among the fT3SS components in the intact *B. burgdorferi* flagellar motor, we constructed a model of the fT3SS machine and its surrounding C-ring complex based on the available homologous structures. We first built the model of the FlhA_C_ nonameric ring based on the homologous structures from other bacteria (Abrusci et al., 2013; Saijo-Hamano et al., 2010) (Fig. 5). The entire ring fits well into the torus-like density (Fig. 5B, C), suggesting that the FlhA complexes also form a nonameric ring in *B. burgdorferi*. Three subdomains (SD1, SD3 and SD4) of FlhA_C_ are located inside the nonameric FlhA_C_ ring, whereas the SD2 domain is located outside of the ring (Fig. S7). The distance between the FlhA_C_ ring and the cytoplasmic membrane is about 6 nm. The FlhA_C_ is linked to the FlhA trans-membrane domain embedded in the cytoplasmic membrane under the MS-ring. The central channel of the export apparatus appears to be aligned with the central axis of the MS-ring and the ATPase complex (Fig. 5D).

**Figure 5.**
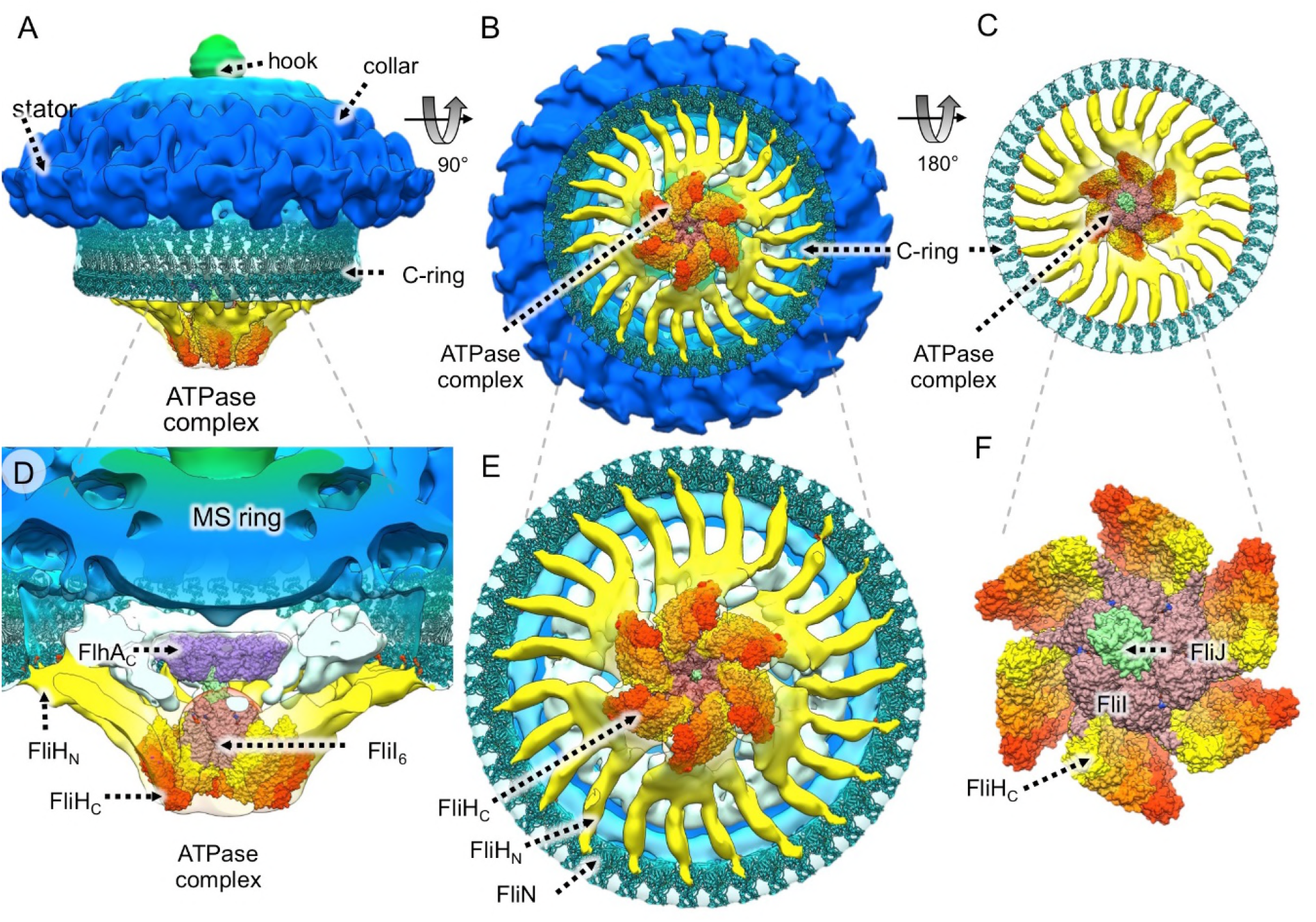
Proposed molecular architecture of the fT3SS in the context of the flagellar motor. Crystallographic structures of FliH, FliI, and FliJ from other organisms (see Materials and Methods) were positioned within the CryoET-derived density map of the large hexametric complex attached to the C-ring protein FliN through FliH links. As there are the 23 FliH ‘spokes’ in the average structure, we speculate there are 46 copies of the FliN tetramer, as well as 46 copies of FliG_MC_-FliM_M_ complex. (A) A side view of the structure of the WT flagellar motor with the assembled C-ring (FliG, FliM and FliN) and an ATPase complex (FliH, FliI and FliJ). (B) A bottom-up view of the C-ring and the ATPase complex. (C) A top-down view of the assembled C-ring and the ATPase complex. (D) A sliced, enlarged view of the ATPase complex and its interactions with FlhA and FliN. (E) An enlarged view of the assembled C-ring and the ATPase complex. The hydrophobic surface (formed by Val-128, Val-129 and Val-130) of FliN interacts with the FliH linker (yellow). (F) A close-up, top-down view of the assembled ATPase complex in which six FliI monomers form the hub and at least 23 FliH dimers form the spoke-like linkers.

The ATPase complex can be divided into two major components: a large central hub, and 23 spoke-like linkers extending to the C-ring (Fig. 5). The ATPase complex was originally proposed to form a hexamer (Claret et al., 2003; Fan and Macnab, 1996; Imada et al., 2007) and is part of the density beneath the FlhA_C_ ring, as suggested by analysis of a Δ*fliI* mutant in *Campylobacter jejuni* (Chen et al., 2011) and in *B. burgdorferi* (Lin et al., 2015). FliI, FliH, and FliJ are known to form a large complex that delivers the chaperone-substrate complex to the export gate (Fraser et al., 2003; Minamino and Macnab, 2000). *B. burgdorferi* contains the homologs of these proteins (Fig. S8). Therefore, we postulate that the hexametric density is composed mainly of the FliI/FliJ complex. Based on its similarity with a portion of the F_0_F_1_ATPase, the FliI/FliJ complex was modeled by aligning the monomer structures of *Salmonella* FliI and FliJ to the α_3_β_3_ and γ parts of F_0_F_1_ATPase, respectively (Ibuki et al., 2011; Imada et al., 2007). The pseudo-atomic structure of FliI/FliJ fits well into the spherical density (Fig. 5C). The density of FliJ is not well resolved in our maps, probably because of its small size or dynamic nature (Ibuki et al., 2011). The N-and C-termini of FliJ insert into the middle of six FliI subunits, while the middle part of FliJ inserts into the nonameric FlhA_C_ ring (Fig. 5C).

A FliH dimer is known to form a stable complex with the FliI ATPase (Minamino and Macnab, 2000). The C-terminal domain of FliH is involved in binding to FliI, while a small central region of FliH is essential for formation of the FliH dimer (Gonzalez-Pedrajo et al., 2002). The N-terminal domain is important for FliH-FliN interactions (Minamino et al., 2009). The C-terminal domain of the FliH dimer interacts with the N-terminal domain of FliI, while the N-terminal domain of the FliH dimer can extend toward FliN at the bottom of the C-ring. We propose that 3 or 4 FliH dimers form each of the hub-like structures. We speculate that one FliH dimer directly binds to one FliI monomer, while others bind to the adjacent FliH dimer in a parallel fashion. In total, there are six bundles of FliH dimers, each of which interacts with one FliI monomer (Fig. 5E, F). The N-terminal domain of the FliH dimer binds to FliN at the bottom of the C-ring (Fig. 5E), and the hydrophobic patch (L85, T110, V128, V130, F135) at the C-terminus of FliN has been reported to interact with FliH (Minamino et al., 2009). The atomic models of FliG, FliM and FliN (Vartanian et al., 2012), which were docked into the C-ring, reveal that these hydrophobic residues of FliN (labeled red in Fig. 5D, Movie S2) are near the tip of the FliH linker.

### Variable conformations of the ATPase complex and the C-ring

The C-ring is thought to rotate together with the MS ring and the flagellar filament, although the rotation of the C-ring has not been directly visualized. The detail of our *in situ* structures is not sufficient to visualize the C-ring and its rotation. However, we are able to resolve multiple FliH linkers between the ATPase complex and the C-ring. To our surprise, classification of the WT motors resolved multiple conformations, in which the ATPase complex apparently adopts different orientations with respect to the collar and the stator (Fig. 6). Specifically, in the four classes shown in Fig. 6, the collar and stator are in a similar orientation, however, the ATPase complexes in classes 03, 05, 08 rotate about 7°, 13°, 20° from class 00, respectively. As the overall model of the C-ring and the ATPase complex fits well into the class averages, we propose that the C-ring and the ATPase can rotate as a large, rigid body (Movie S3).

**Figure 6.**
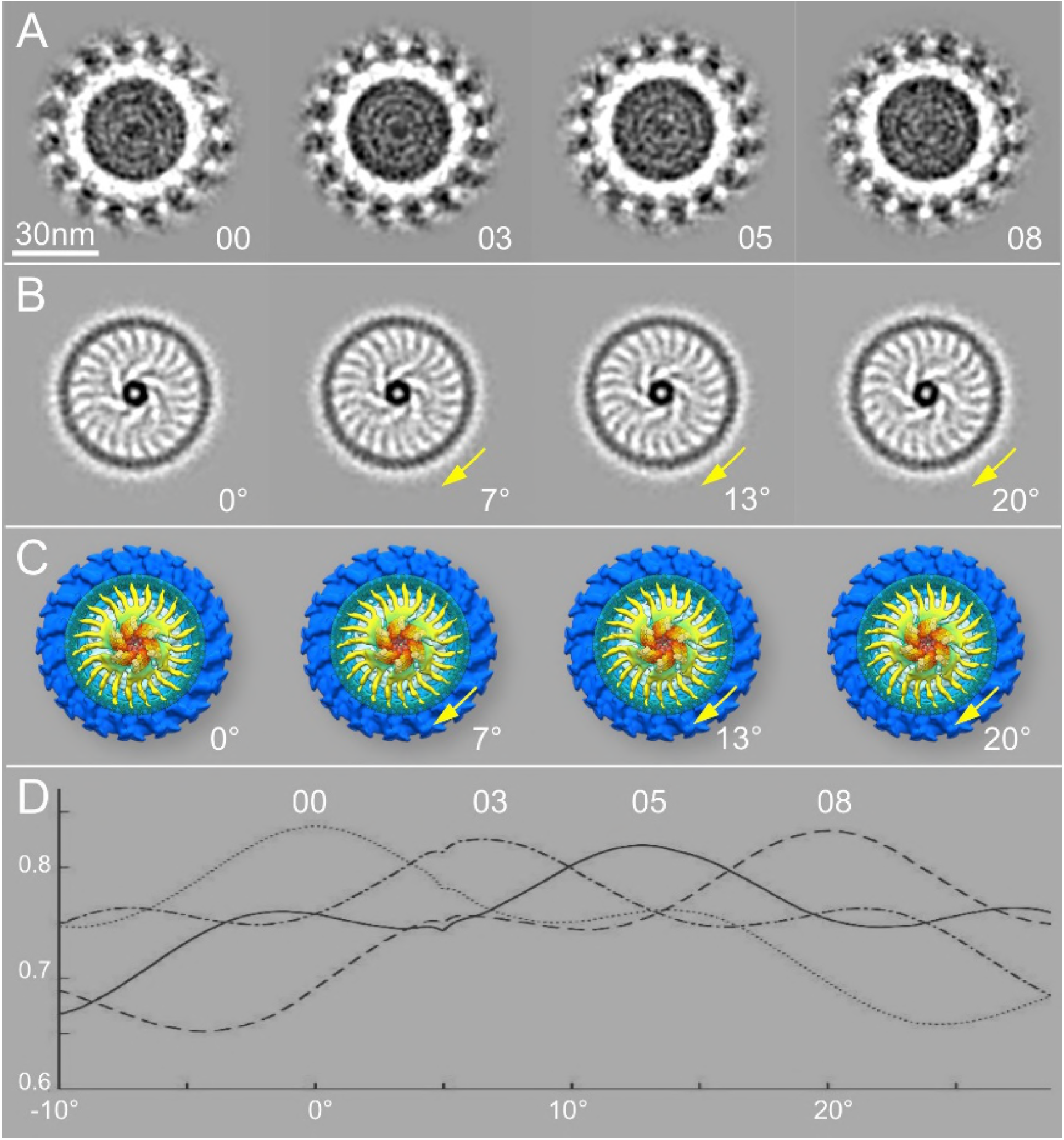
Variable conformations of the ATPase complex and the C-ring. (A) Sections of four class averages at the level of the 16 circumferential stator densities. Note that the stator densities exhibit very similar patterns. (B) Sections of the same class averages shown in panel A, but taken at the level of the FliI/FliH assembly and the C-ring. The sections show the ATPase complex in slightly different orientations. There are different rotations in classes 03, 05, and 08 relative to class 00. (C) Cytoplasmic views of the ATPase complexes from the four class averages, corresponding to the cross sections in panel B, respectively. (D) Cross correlation coefficient (CCC) between class averages. Note that the peak of the CCC for class 00 happens at 0° (without any in-plane rotation). The CCC peak for class 03 is located at ~7°, the CCC peak for class 05 at ~13°, and the CCC peak for class 08 at 20°.

## DISCUSSION

T3SSs in bacterial flagella and injectisomes are highly conserved. The flagella are elaborate self-assembling machines that serve as the main organelles for bacterial motility. The injectisomes are specialized nanomachines deployed by many important human pathogens such as *Salmonella* spp., *Shigella* spp. and *Pseudomonas* to deliver virulence effectors into eukaryotic cells. Our previous studies revealed key intermediates of fT3SS-mediated assembly in *B. burgdorferi* (Zhao et al., 2013) and overall architectures of the vT3SS machines in *Shigella* and *Salmonella* (Hu et al., 2017; Hu et al., 2015). Here we focus on the structure and function of the fT3SS machine in periplasmic flagella by systematically analyzing mutants lacking key fT3SS components and comparing them to the vT3SS machines (Fig. 7). The overall organization of the fT3SS machine in the *B. burgdorferi* periplasmic flagella shares many similar features observed with the fT3SS machine in the *E. coli* external flagella (Zhu et al., 2017) and the vT3SS machines in *Shigella* and *Salmonella* (Hu et al., 2017; Hu et al., 2015; Kawamoto et al., 2013). However, there are considerable differences between the fT3SS and vT3SS.

**Figure 7.**
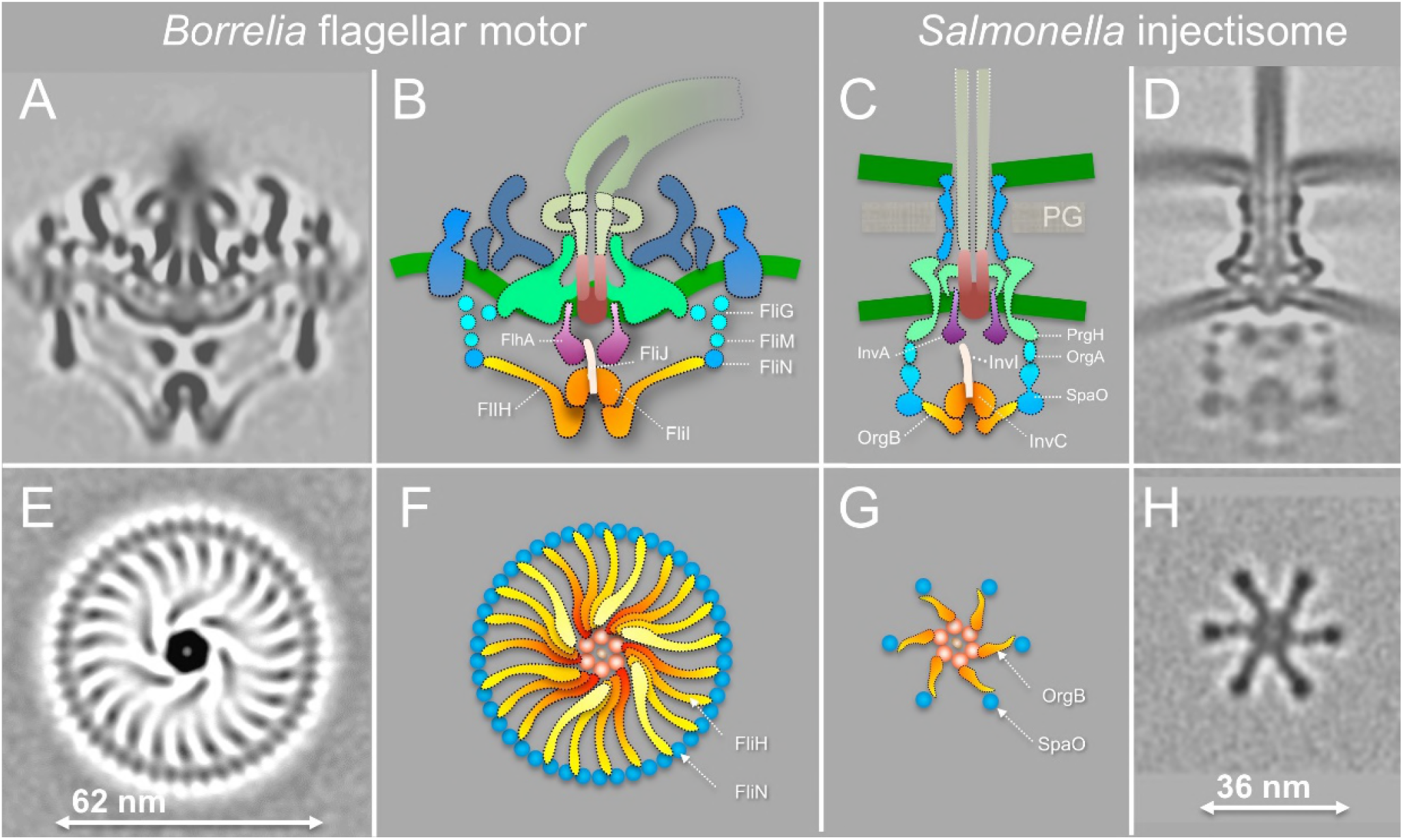
Comparison of the fT3SS from *B. burgdorferi* and the vT3SS from *Salmonella*. (A) A central section from the WT *B. burgdorferi* motor. (B) The fT3SS in the spirochete motor consists of the ATPase complex (orange) and the export apparatus (purple) underneath the MS-ring. (C, D) The vT3SS from *Salmonella* injectisome is modeled in a similar color scheme (Hu et al., 2017; Hu et al., 2015). The difference between the two T3SSs is striking in a comparison of the cross sections of their ATPase complexes. Note that the C-ring from the *B. burgdorferi* motor is a continuous ring with ~46 copies of FliN tetramer. There are 23 visible FliH linkers (E, F). There are six pods in *Salmonella* injectisome. Only six linkers of the FliH homolog OrgB connect the ATPase complex to the SpaO molecules that compose the pod of the injectisome.

Five conserved membrane proteins (FlhA, FlhB, FliP, FliQ, and FliR) form the export apparatus beneath the MS-ring in *B. burgdorferi*. They are essential for the export of flagellar proteins and for motility. FliP, FliQ, and FliR likely form an export gate in the cytoplasmic membrane. FlhA forms a nonameric ring complex, which provides a docking site for the ATPase complex and substrates. The overall structure of the export apparatus in the *B. burgdorferi* flagellar motor is similar to that in the *Salmonella* injectisome (Fig. 7) and external flagella. Presumably, similar mechanisms are utilized for substrate export.

The ATPase complex of the *B. burgdorferi* periplasmic flagella is quite different from that in the *Salmonella* injectisome (Fig. 7) and the *E. coli / Salmonella* external flagellum (Kawamoto et al., 2013; Zhu et al., 2017). The ATPase complex is surrounded by a large continuous C-ring of the *B. burgdorferi* flagellar motor, while it is linked to six “pods” in the *Salmonella* injectisome (Compare Fig. 7E, F with G, H). In particular, we observed spoke-like linkers between the ATPase and the C-ring for the first time in any bacterium, as they are not observed in the recent structures from the external flagella (Kawamoto et al., 2013; Zhu et al., 2017). Surprisingly, there are about 23 linkers (Fig. 7E, F). They are considerably longer (6 nm vs. 3 nm) in the fT3SS machine than in the vT3SS machine, mainly because the C-ring is much larger than the pod array (62 nm vs. 36nm in diameter) (Fig. 7). Previous studies provided evidence that OrgB (a FliH homolog) forms the spoke-like structure and interacts with the ATPase complex and SpaO (a FliN homolog) of the vT3SS machine (Hu et al., 2017). In the *B. burgdorferi* flagellar motor, the linker between the ATPase and C-ring is likely formed by multiple FliH molecules. Consistent with the longer linker region, FliH of *B. burgdorferi* is significantly larger (305 residues) than its homolog OrgB (170 residues). Thus, the ATPase complex not only provides a large docking platform for substrates recruitment and secretion, but also supports the integrity of the C-ring, which undergoes rotation and switches between clockwise and counterclockwise rotation.

We observed many different orientations of the ATPase complex relative to the periplasmic structures of the motor, suggesting that the C-ring and the ATPase complex rotate together beneath the MS-ring. The rotation of the C-ring is driven by sixteen stators that surround the C-ring and a spirochete-specific periplasmic collar (Moon et al., 2016). In contrast, the pods found in *Salmonella* injectisomes do not appear to rotate, although OrgB and SpaO likely undergo high turnover with a cytoplasmic pool. These key differences between the fT3SS and vT3SS underline the distinct mechanisms involved in the assembly and function of flagella and injectisomes.

The assembly of the flagellum can be divided into two distinct processes. The first stage includes the formation of the MS-ring, C-ring, export apparatus, and the stator. The second stage, which includes the assembly of the rod, hook, and filament, is mediated by the fT3SS. It is generally thought that the MS-ring is the first unit assembled and is central to flagellar assembly and function (Kubori et al., 1992). Recently, fluorescence microscopy was used to investigate dynamic protein exchange in the assembled *E. coli* motor structure. This study suggested that flagellar assembly is initiated by oligomerization of the export protein FlhA, which is followed by the recruitment of the MS-ring protein FliF (Li and Sourjik, 2011). In another study, FlhA localization in *S. enterica* required FliF, FliG, FliO, FliP, FliQ and FliR, suggesting that FlhA assembles into the export gate along with other membrane components in a coordinated manner during the MS-ring formation (Bai et al., 2014). In *B. burgdorferi*, FlhA plays little role in initiation of motor formation, because deletion of *flhA* has no impact on the number of motors per cell tip (Fig. 4F). In contrast, other membrane export proteins (FliP, FliQ, FliR, and FlhB) have considerable impact on motor formation. Even in the absence of FliP, FliQ, FliR, FlhA and FlhB, motor assembly still occurs with very low efficiency. Taken together, our results imply that most export proteins are involved in the coordinated assembly of the MS-ring and export apparatus, whereas FlhA is not critical in this process, at least, in *B. burgdorferi*. Interestingly, similar results were obtained in the *Salmonella* injectisome (Wagner et al., 2010), suggesting that coordination in the assembly of the basal body and export apparatus might be shared by flagella and injectisomes.

In conclusion, our study reveals unprecedented details about the fT3SS machine in the Lyme disease spirochete *B. burgdorferi*. We systematically characterize the fT3SS machine, map the key components, and document their roles in flagellar structure and function. We present the first structural evidence that the distinct ATPase complex of the fT3SS machine is attached to the flagellar C-ring through multiple spoke-like linkers comprised of FliH. The novel architecture not only strengthens the C-ring, but also enables an optimal translocation of substrates through the ATPase complex and the export apparatus. Remarkably, the ATPase complex together with the C-ring can adopt variable orientations, implying that the fT3SS machine undergoes rotation in concert with the flagellar C-ring. Therefore, our studies not only provide a structural framework for a better understanding of the fT3SSs, but also underscore the striking differences between flagella and their evolutionally related bacterial injectisomes.

## ACKNOWLEDGMENTS

We thank Drs. William Margolin, Michael Manson, James Stoops, and Shenping Wu for suggestions and comments. We thank Patricia Rosa for sharing reagents. This work was supported by grants from National Institute of Allergy and Infectious Diseases (NIAID) (R01AI087946, R01AI078958, R01DE023080, and R21AI113014), National Institute of Arthritis and Musculoskeletal and Skin Diseases (NIAMS) (R01AR060834), and Welch Foundation (AU-1714).

## EXPERIMENTAL PROCEDURES

### Bacterial strains and growth conditions

High-passage *Borrelia burgdorferi* strain B31A (WT) and its isogenic mutants (Table S2) were grown in BSK-II liquid medium supplemented with 6% rabbit serum or on semi-solid agar plates at 35°C in the presence of 2.5% carbon dioxide as previously described (52, 53).

### Construction of deletion mutants of *fliP, fliQ, fliR, flhA, flhB*, and *FliO*

All mutants studied in this communication were constructed at East Carolina University. Single mutants were constructed using a gene inactivation methodology that creates deletion mutants without imposing any polar effects (Motaleb et al., 2011). The *fliP* gene (gene locus *BB0275*; a 765 bp gene) was inactivated by replacing *fliP* with the *aadA* coding sequence by using overlapping PCR, as schematically shown in Figure S10. PCR was used to amplify three regions of DNA in three steps. In step one, each DNA region was amplified separately by using PCR pairs P1-P2 (5’-upstream *fliP*, 3’-upstream *fliP*), P3-P4 (*aadA* coding sequence of streptomycin resistance gene), and P5-P6 (5’-downstream *fliP*, 3’-downstream *fliP*). Primers P2, P3, P4, and P5 (Table S3) contain several overlapping base pairs, as indicated by different colors in Figure S10. In step two, a PCR product was obtained by using primers P1 and P4 and the purified DNA products for upstream *fliP* and *aadA* as templates. In step three, the final PCR product was obtained by using primers P1 and P6, and the purified DNA products of upstream *fliP-aadA* and downstream *fliP* as a template to amplify the upstream *fliP-aadA*-downstream *fliP* DNA construct. The final PCR product yielded a 2,462-bp product that was gel purified and cloned into the pGEM-T Easy vector (Promega Inc.), and then confirmed by PCR and restriction mapping. PCR-amplified DNA was electroporated into B31A competent cells and plated in BSK-II medium containing 100 μg / ml streptomycin (Motaleb et al., 2007). Construction of the *fliQ, fliR, flhB*, and *flhA* mutants were similarly achieved. Construction of the *FliO* mutant was achieved by replacing the respectable gene with the *aph1* coding sequence for kanamycin resistance, as described above. Resistant clones were analyzed by PCR for the confirmation of homologous recombination (Figure S11). Primers used in creating these mutants are listed in Table S3.

### Construction of point mutants *flhA* D158E and *flhA* D158N in *B. burgdorferi*

Wild type *B. burgdorferi flgB* operon contains 26 genes including the target *flhA* (Fig. S10B, top panel), which is transcribed by σ^70^. The point mutants were created as follows. Using WT *B. burgdorferi* B31A cells DNA as the template, the left arm (2094 bp of*flhA*) and right arm (1060 bp downstream of*flhA*) were PCR amplified. Using overlapping PCR, a previously described promoter-less kanamycin cassette (*Pl-Kan*; 846 bp) was inserted between the left arm and right arm, as depicted in Figure S10B, and subsequently cloned into pGEM-T Easy vector yielding pGEM-T Easy*::flhA-Pl-Kan*. Using pGEMTeasy::*flhA-Pl-Kan* as the template, point mutations D158E (GAT to GAA) and D158N (GAT to AAT) were made by QuikChange II Site-Directed Mutagenesis Kit (Agilent Technologies Inc.), yilding pGEM-T Easy::*flhAD158E-Pl-Kan* and pGEM-T Easy::*flhAD158N-Pl-Kan*, respectively. The linearized DNA were then electroporated separately into the competent *B. burgdorferi* B31A cells as described above, and transformants were selected with 200 μg/ml Kanamycin. The antibiotic resistant colonies were sequenced to confirm the point mutations.

### Construction of a quintuple mutant Δ*fliP-flhA*

Deletion of the *fliP, fliQ, fliR, flhB, and flhA* genes was achieved by utilizing the Cre-lox recombination system (Bestor et al., 2010) (Figure S12). Briefly, *LoxP* sites were introduced into *fliP* and *flhA* genes in the same orientation and chromosome causing the deletion of the sequence containing genes *fliP, fliQ, fliR, flhB*, and *flhA. LoxP* sites were introduced by PCR amplifying *fliP* and *flhA*, containing a single HindIII site, using PCR primer pairs P31-P32 and P33-P34 (Table S3). The resultant PCR products were gel purified and cloned into the pGEM-T Easy vector (Promega). A HindIII restriction site was engineered flanking the *loxP* site with streptomycin resistance (pABA07) by PCR primer pairs P35-36 and cloned into the pGEM-T Easy vector. The *loxP* site with streptomycin resistance cassette in pGEM-T Easy and *loxP* site with kanamycin resistance cassette (pABA14) were digested with HindIII and cloned into plasmids containing the *fliP* and *flhA* genes in pGEM-T Easy, respectively, which were also digested with HindIII, to create the *loxP* insertion mutant vectors. The integrity of the *loxP* insertion mutant vectors and orientation of the *loxP* sites were confirmed by PCR and restriction mapping (Figure S11).

### Determination of polar effect on downstream gene expression

Our novel gene inactivation system does not impose any polar effects, as we have confirmed and verified previously (Motaleb et al., 2011; Zhao et al., 2013). However, we still determined the effect of a deletion mutant on the expression of the downstream genes using qRT-PCR as described previously (Sze et al., 2013). Total RNA was extracted from exponentially-grown *B. burgdorferi* (10 ml) cells by using Direct-zol™ RNA MiniPrep Kit (Zymo Research). To ensure that the samples were free of contaminating genomic DNA, the RNA preparation was digested with Turbo DNase I (Ambion) overnight. The concentration and purity of each RNA sample were measured via spectrophotometry (ND-1000 spectrophotometer; NanoDrop Technologies, Inc., Wilmington, DE) and were also assessed by gel electrophoresis. Samples were checked for contamination of genomic DNA by PCR, using *B. burgdorferi* enolase primers. First-strand cDNA was prepared by using AffinityScript cDNA Synthesis Kit (Agilent) according to the manufacturer’s instructions. The resulting cDNA was amplified using a CFX96 RealTime System (Bio-Rad), with a final reaction volume of 25 μl that contained 10 ng of cDNA, Power SYBR^®^ Green PCR Master Mix (Life Technologies), and *B. burgdorferi* gene-specific primers. *B. burgdorferi* enolase was used as an internal control. Real-time PCRs were carried out in triplicate, with consistent results (Figure S13, S14).

### Dark-field Microscopy to determine motility and bacterial morphology

Live *B. burgdorferi* cells were observed under a dark-field microscope (Zeiss Axio Imager. M1) connected to an AxioCam digital camera. Exponentially growing cells were examined for their shape and motility. Almost all mutants were non-motile and rod-shaped (see Fig. S3 as an example).

### Frozen-hydrated EM sample preparation

The frozen-hydrated specimens were prepared as previously described (Liu et al., 2009). Briefly, *B. burgdorferi* culture was centrifuged at 5,000 × g for 5 minutes. The pellet was suspended with 1.0 ml PBS. The cells were centrifuged again and suspended in 50~80 μl PBS. The cultures were mixed with 10 nm colloidal gold and were then deposited onto freshly glow-discharged, holey carbon grids for 1 min. Grids were blotted with filter paper and then rapidly frozen in liquid ethane, using a homemade gravity-driven plunger apparatus.

### Cryo-electron tomography

Frozen-hydrated specimens were imaged at −170 °C using a Polara G2 electron microscope (FEI) equipped with a field emission gun and a 16 megapixel CCD camera (TVIPS). The microscope was operated at 300 kV with a magnification of 31,000 ×, resulting in an effective pixel size of 5.7 Å after 2×2 binning. Using the FEI “batch tomography” program, low-dose, single-axis tilt series were collected from each cell at −6 to −8 μm defocus with a cumulative dose of ~100 e^-^/Å^2^ distributed over 87 images and covering an angular range of −64° to +64°, with an angular increment of 1.5°. SerialEM was recently used to collect tilt series from WT cells at a Gatan K2 Summit direct detector device (DDD) with dose fractionation mode. The microscope was operated at a magnification of 9,400 ×, resulting in an effective pixel size of 4.45 Å without binning and a cumulative dose of ~60 e^-^/Å^2^ distributed over 61 stacks. Each stack contains 8 images. We developed Tomoauto (a wrapper library) to facilitate the automation of cryo-ET data processing (Hu et al., 2015). The main executable encompasses: drift correction of dose-fractionated data using motioncorr (Li et al., 2013) and the assembly of corrected sums into tilt-series; alignment of tilt-series by IMOD (Kremer et al., 1996); reconstruction of tilt-series into tomograms by TOMO3D (Agulleiro and Fernandez, 2011).

### 3-D image processing and sub-tomogram averaging

In total, 2,846 tomographic reconstructions of Δ*flhA*, Δ*flhB*, Δ*FliO*, Δ*fliP*, Δ*fliQ*, Δ*fliR*, Δ*fliP-flhA* and WT cells were generated and 12,658 flagellar motor sub-tomograms (256×256×256 voxels) were extracted (Table S4). The sub-tomogram analysis was utilized as previously described (Liu et al., 2009; Zhao et al., 2013). Briefly, the initial orientation of each motor was estimated by the center coordinates of the flagellar C-ring and the collar, thereby providing two of the three Euler angles. To accelerate image analysis, 4×4×4 binned sub-tomograms (64×64×64 voxels) were used for initial alignment. Then, the original sub-tomograms (256×256×256 voxels) were utilized for further image analysis. Multivariate statistical analysis and hierarchical ascendant classification were then applied to analyze the intact motor (*Liu et al., 2010b; Winkler, 2007; Winkler et al., 2009*). Relevant voxels of the aligned sub-volumes were selected by specifying a binary mask of the motor. Class averages were computed in Fourier space, so the missing wedge problem of tomography was minimized. All class averages were further aligned with each other to minimize differences in motor orientation. Because of the predominant 16-fold-symmetric feature of the flagellar motor, the structure of the T3S machine is not well resolved. A novel procedure was developed in which specific substructures of interest are classified and aligned without applying rotational symmetry. Specifically, classification focusing on the export apparatus revealed significant details in its overall structure and interaction with the C-ring. The cytoplasmic portion of the export apparatus complex shows evident features in 6-fold symmetry, while the periplasmic features maintain in 16-fold symmetry (Fig. 2G).

The average structure of the spoke from WT was generated by aligning the ATPase region and classification on the spoke region using eigenimages 1 to 18 (See Fig. S1 A). The first 40 eigenimages of the data set of ~30,000 sub-tomograms show different symmetry of the spoke region (See Fig. S1B). Eigenimages 01 and 02 exhibit 23-fold symmetry. Eigenimages 04 and 07 exhibit 22-fold symmetry. Eigenimage 08 and 09 exhibit 21-fold symmetry. Eigenimage 10 and 11 exhibit 24-fold symmetry. The eigenimages were ranked by the highest values, which accounts for larger percentage of the total variance of the data set.

To obtain the symmetry mismatching structures in Fig 6. and compare the rotation angles, the class averages shows the spoke-like links were selected and aligned based on the region of the C-ring and the ATPase with links. Then they were classified on the collar and stator region. The new class averages that show the symmetry of the collar and stator were selected and aligned by the collar and stator region with spin alignment only. The spin rotation angle were recorded and compared.

### 3-D visualization and modeling

UCSF Chimera (Pettersen et al., 2004) was used for 3-D visualization of flagellar motors. Using “match maker” in UCSF Chimera, we built the nonameric ring model of FlhA_C_ based on the homologue MxiA_C_ (PDB: 4A5P) from *S. flexneri* (Abrusci et al., 2012). The model of the FliI-FliJ complex was built by aligning the hexameric FliI ring (PDB: 2DPY) (Imada et al., 2007) and the monomer FliJ (PDB: 3AJW) (Ibuki et al., 2011) with α, β and γ subunit of bovine F_1_ ATPase (PDB: 1E79) (Gibbons et al., 2000) (Fig. S9). The models are then docked into 3-D density maps by using the function “fit in map” in UCSF Chimera (Pettersen et al., 2004) (Figure 5). FliN is organized in doughnut-shaped tetramers (Paul and Blair, 2006). Together with a recent crystal structure FliM_M_-FliG_MC_ complex from *Thermotoga maritima* (PDB: 4FHR) (Vartanian et al., 2012), the FliN-FliM_M_-FliG_MC_ complexs fit well into the bulge density at the bottom of the C-ring (Figure 5G-I; Video 1). As V111, V112, V113 *(E. coli)* are in the hydrophobic patch and interaction with FliH (Paul et al., 2006), we speculate those three Valine facing towards the FliH link. Those three Valine correspond to V128, V129 and V130 in *T. maritima* (Paul et al., 2006). As a result, when we fit the FliN tetramer ring, we have V128, V129 and V130 (See Movie S2 shown in red) facing towards the FliH link.

---

**Figure S1.**
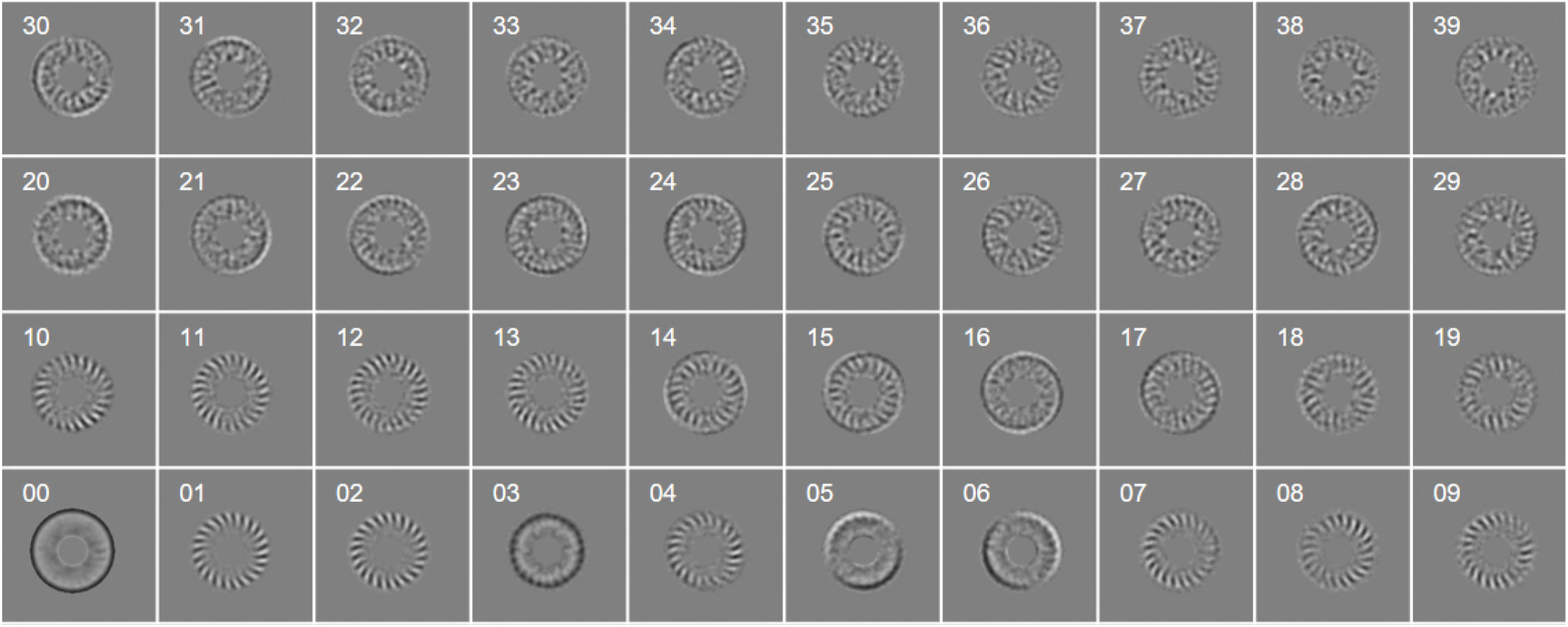
Main eigen images for the classification on the spoke region. The first 40 eigenimages of the data set of ~30,000 sub-tomograms show different symmetry of the spoke region. Those subtomograms were from c6 rotating of ~5000 motors along the axle, as the ATPase domain has the 6fold rotational symmetry. Eigenimages 01 and 02 exhibit 23-fold symmetry. Eigenimages 04 and 07 exhibit 22-fold symmetry. Eigenimages 08 and 09 exhibit 21-fold symmetry. Eigenimages 10 and 11 exhibit 24-fold symmetry.

**Figure S2.**
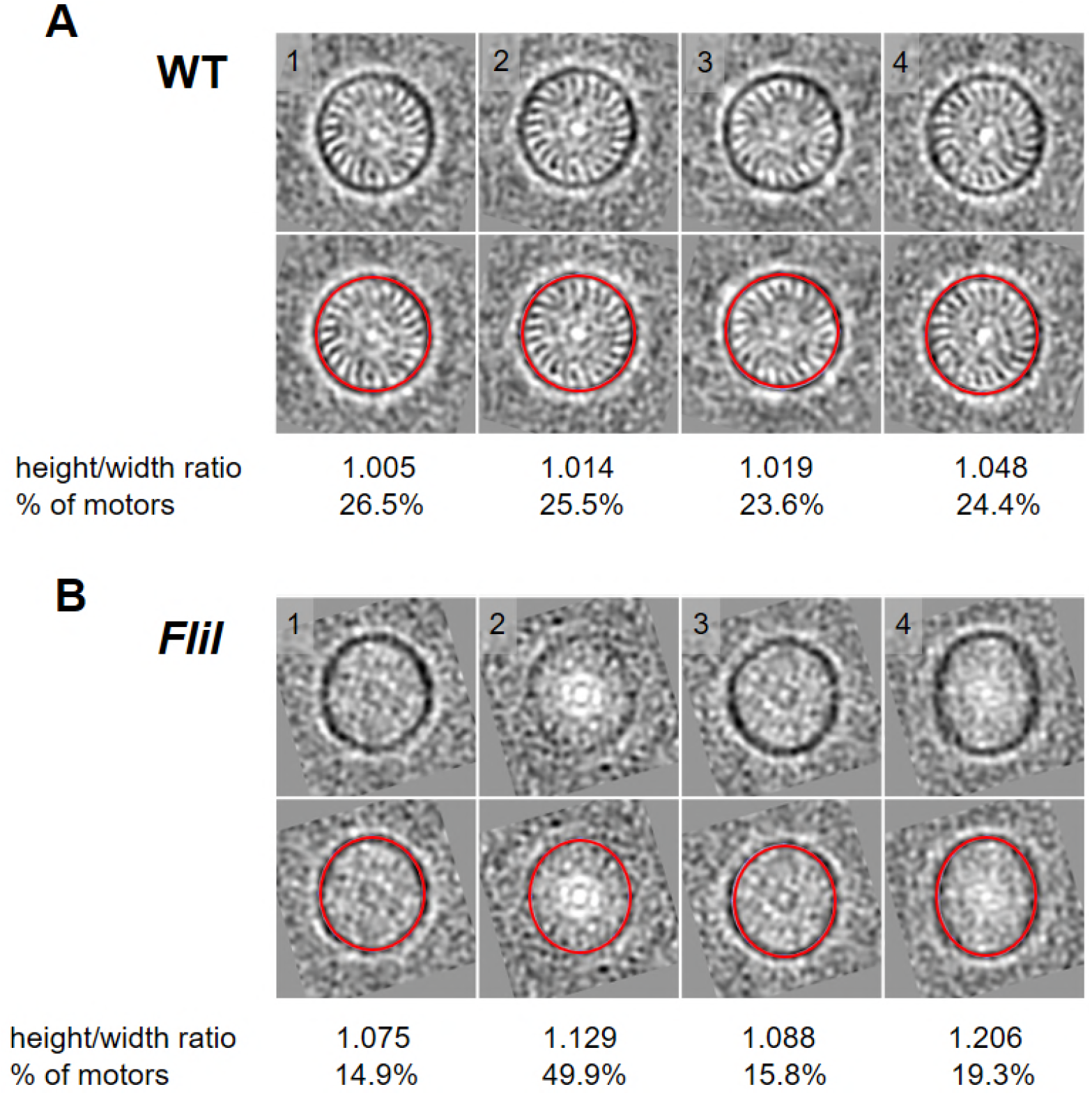
Comparison of flagellar C-ring in the WT and the Δ*fliI* mutant. *B. burgdorferi* flagellar motors from WT and *fliI* mutant were aligned and classified on the C-ring. The class averages were viewed in the cross section of the C-ring (sliced at position C as shown in Figure 2A). (A) Top: cross sections of 4 class averages from wild-type. Bottom: The red circle superimposed on the C-ring measures the height/width ratio of each class average. The ratio and the percentage of motors in each class are shown below the class averages. (B) Top: cross section of 4 class averages from the *fliI* mutant. Bottom: The red circle superimposed on the C-ring measures the height/width ratio of each class average. The ratio and the percentage of motors in each class are shown below the class averages.

**Figure S3.**
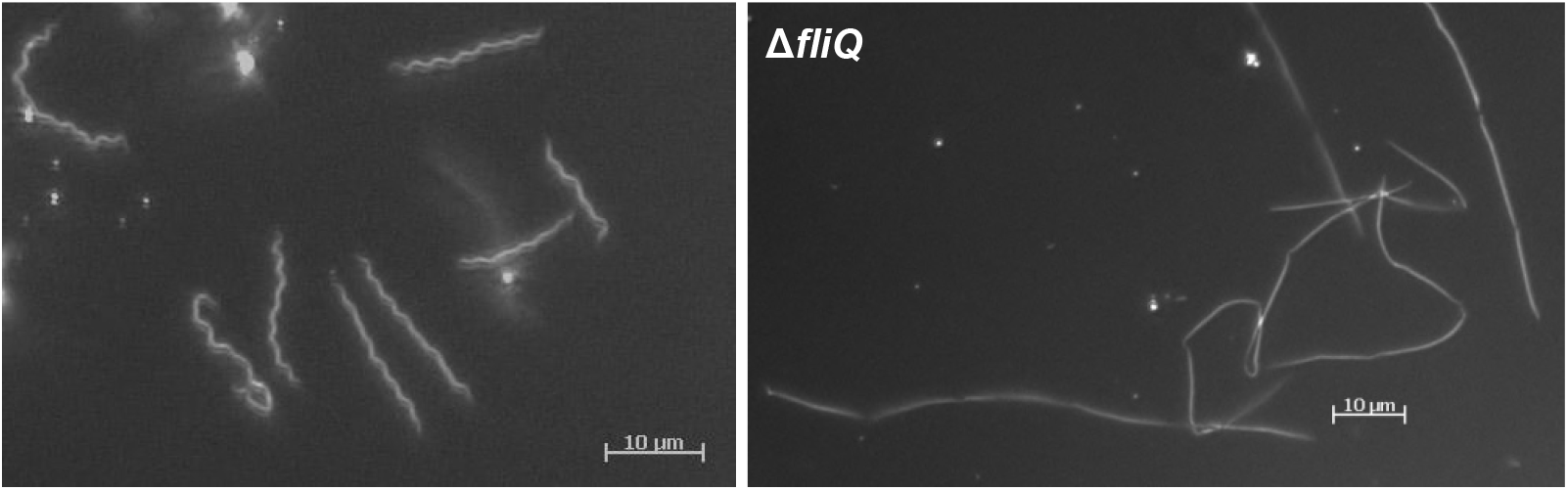
Dark-field images of the generated mutants. A representative dark-field image of motile, flat-wave wild-type (left), and a typical image of the rod-shaped, non-motile mutant *(ΔfliQ)*.

**Figure S4.**
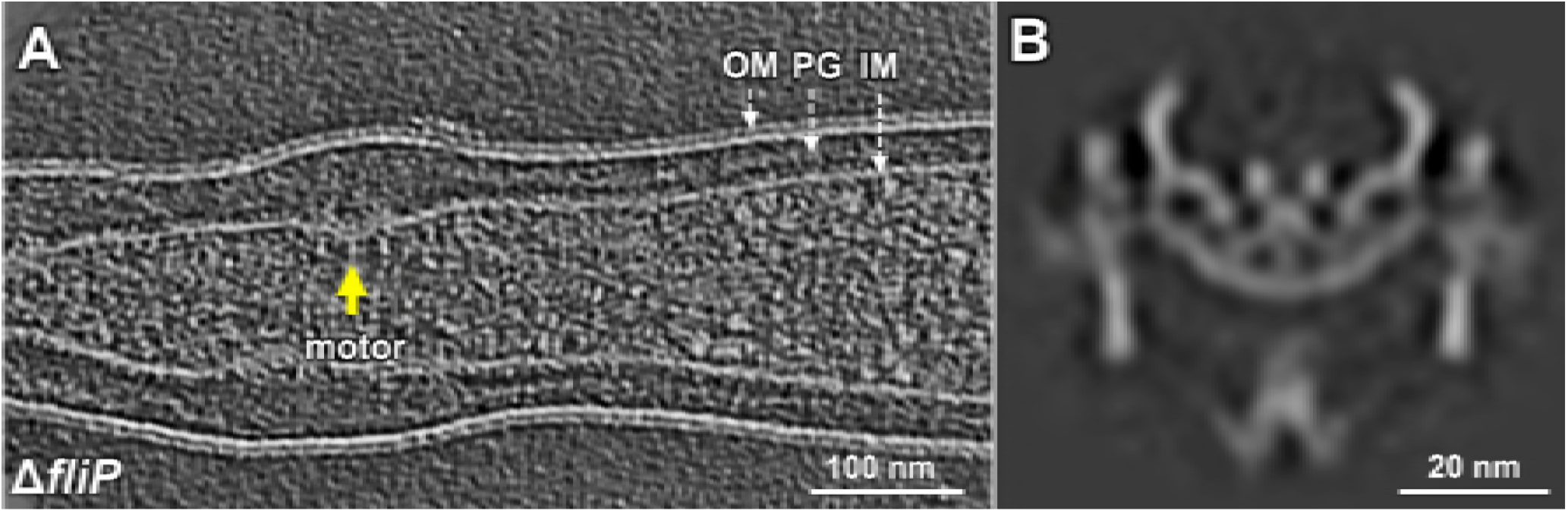
Structural characterization of the flagellar motor in *ΔfliP* cells. (A) A section from a tomogram of *ΔfliP* cells. One motor is highlighted in yellow arrow. (B) A central section of the flagellar motor in *ΔfliP* cells.

**Figure S5.**
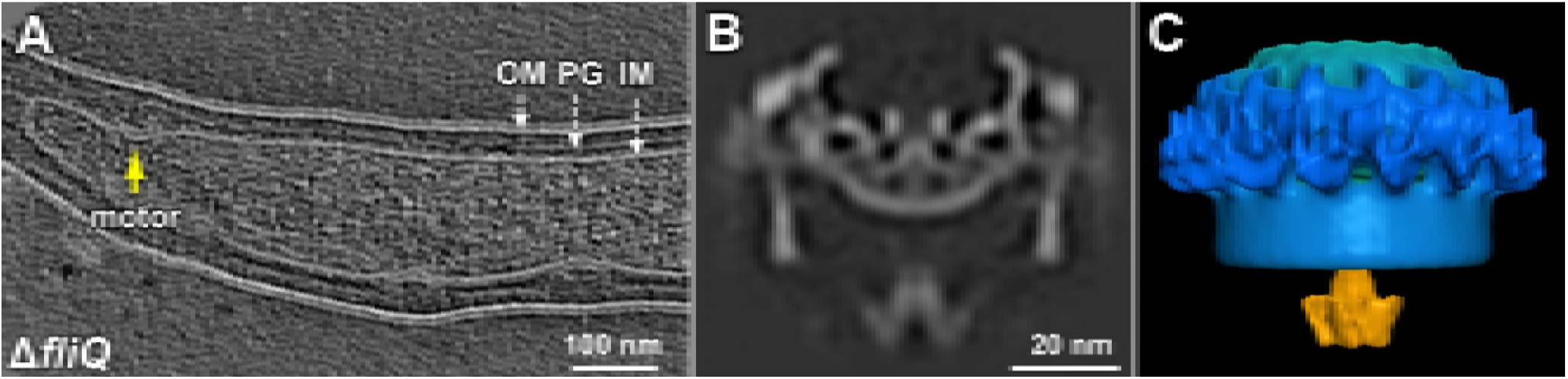
Structural characterization of the flagellar motor in *ΔfliQ* cells. (A) A section from a tomogram of mutant *ΔfliQ* cells. One motor is shown in yellow arrow. (B, C) A central section and a surface rendering of the sub-tomogram average of the flagellar motor in *ΔfliQ* cells, respectively.

**Figure S6.**
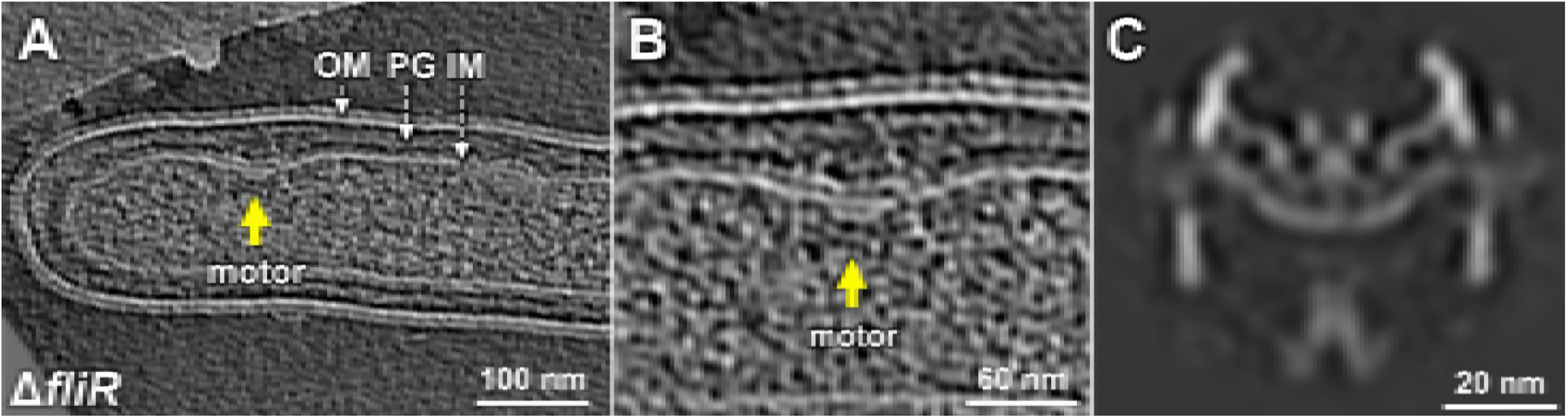
Structural characterization of the flagellar motor in *ΔfliR* cells. (A) A section from a tomogram of *ΔfliR* cells. The yellow arrow indicates one motor. (B) A zoom-in view of the motor. (C) A central section of the sub-tomogram average of the flagellar motor in *ΔfliR* cells.

**Figure S7.**
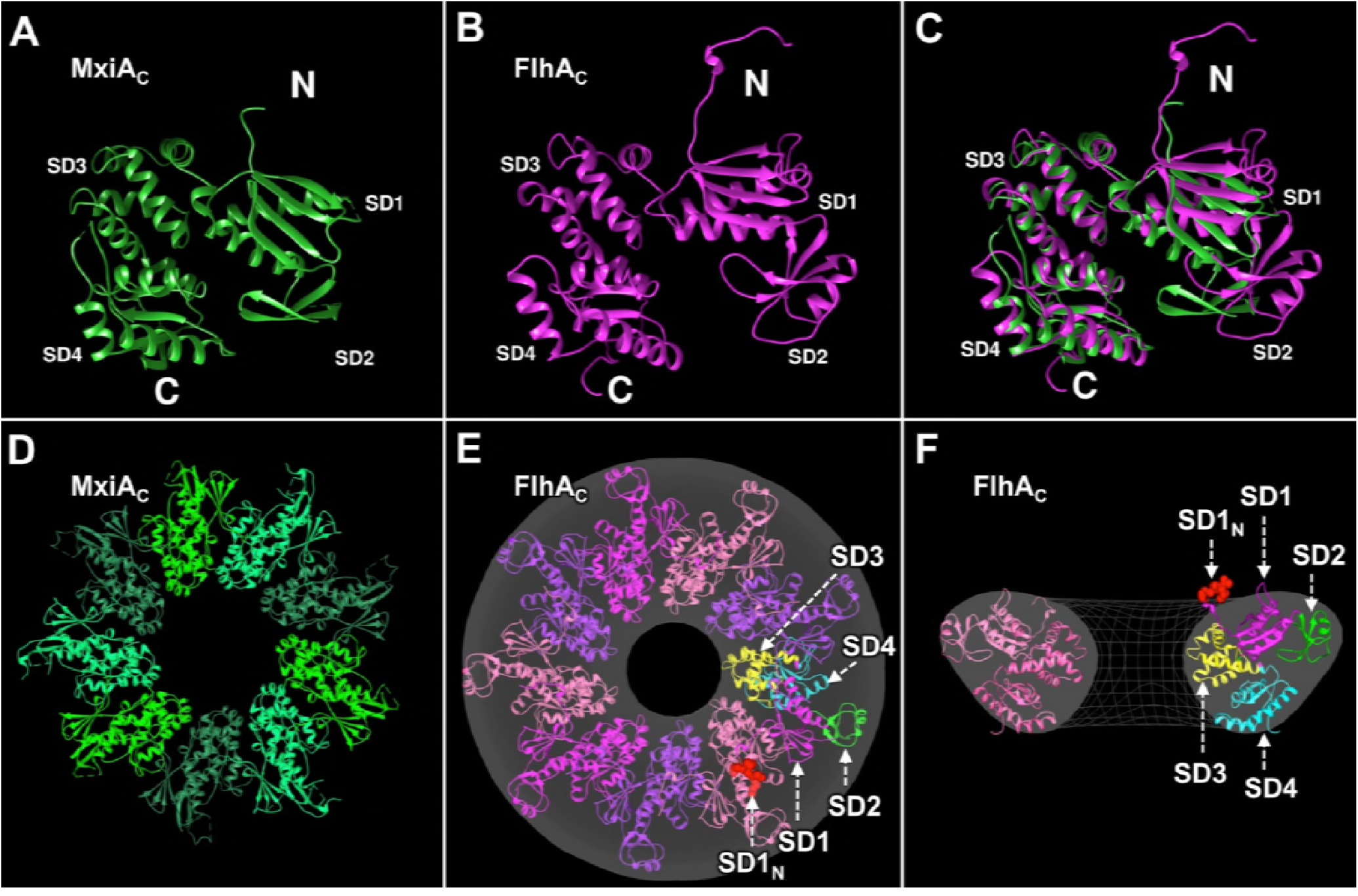
Molecular model of FlhA_c_ nonameric ring. Monomeric structure of the cytoplasmic domain (MxiA _C_) of *Shigella flexneri* MxiA. There are four subdomains (SD1, SD2, SD3, and SD4). (B) Monomeric structure of the cytoplasmic domain (FlhA_C_). Two structures are aligned. (D) The crystal structure of the nonameric ring of MxiA_C_. (E) A model of the nonameric ring of FlhA_C_ was built based on the nonameric ring of MxiA_C_. It fits well into the EM structure. Four subdomains are highlighted in different colors and the N-terminal five residues (SD1_N_) are highlighted in red. (F) A central section of (E).

**Figure S8.**
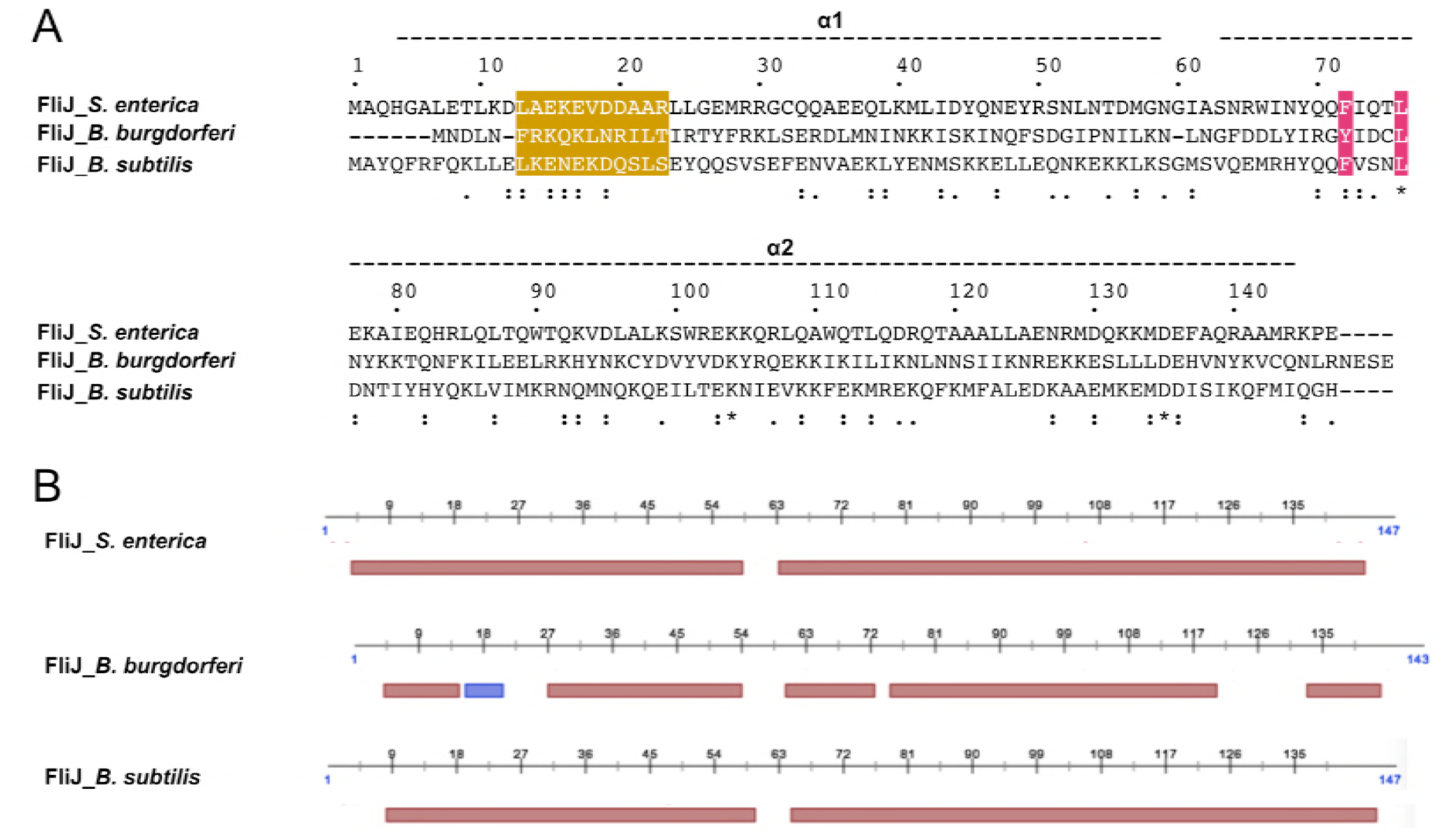
FliJ in *B. burgdorferi* is a homologue of FliJ in other bacteria. (A) Sequences of *S. enterica* FliJ, *B. burgdorferi* FliJ and *B. subtilis* FliJ were aligned using CLUSTALW2. Asterisks under the sequences indicate the identical residues among the three bacteria; dots indicate similar residues. Residues 72 and 76 highlighted by red boxes with white letters were reported to interact with FlhA (Ibuki et al., 2013). Residues 13-24 highlighted by a yellow box with white letters were reported to interact with FliI (Minamino et al., 2011). (B) Secondary structure prediction indicates that FliJ is mainly composed of alpha helices, similar to FliJ.

**Figure S9.**
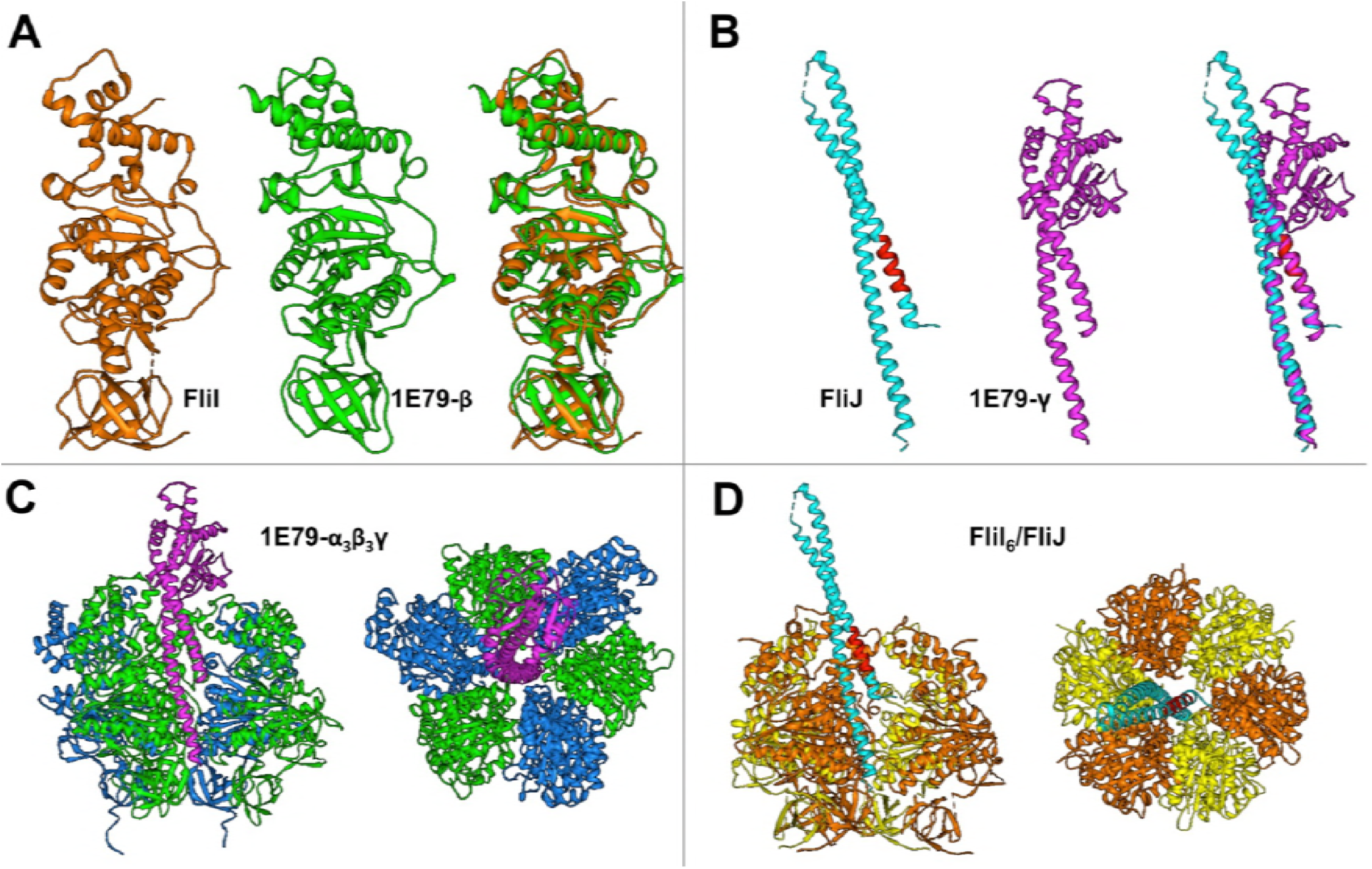
Molecular model of FliI/FliJ complex. (A) The crystal structure of FliI from *S. enterica* is compared with the crystal structure of β subunit of F_1_-ATPase. (B) The crystal structure of FliJ from *S. enterica* is compared with γ subunit of F_1_-ATPase. (C) The crystal structure of F_1_-ATPase (D) A homolog model of FliI / FliJ complex.

**Figure S10.**
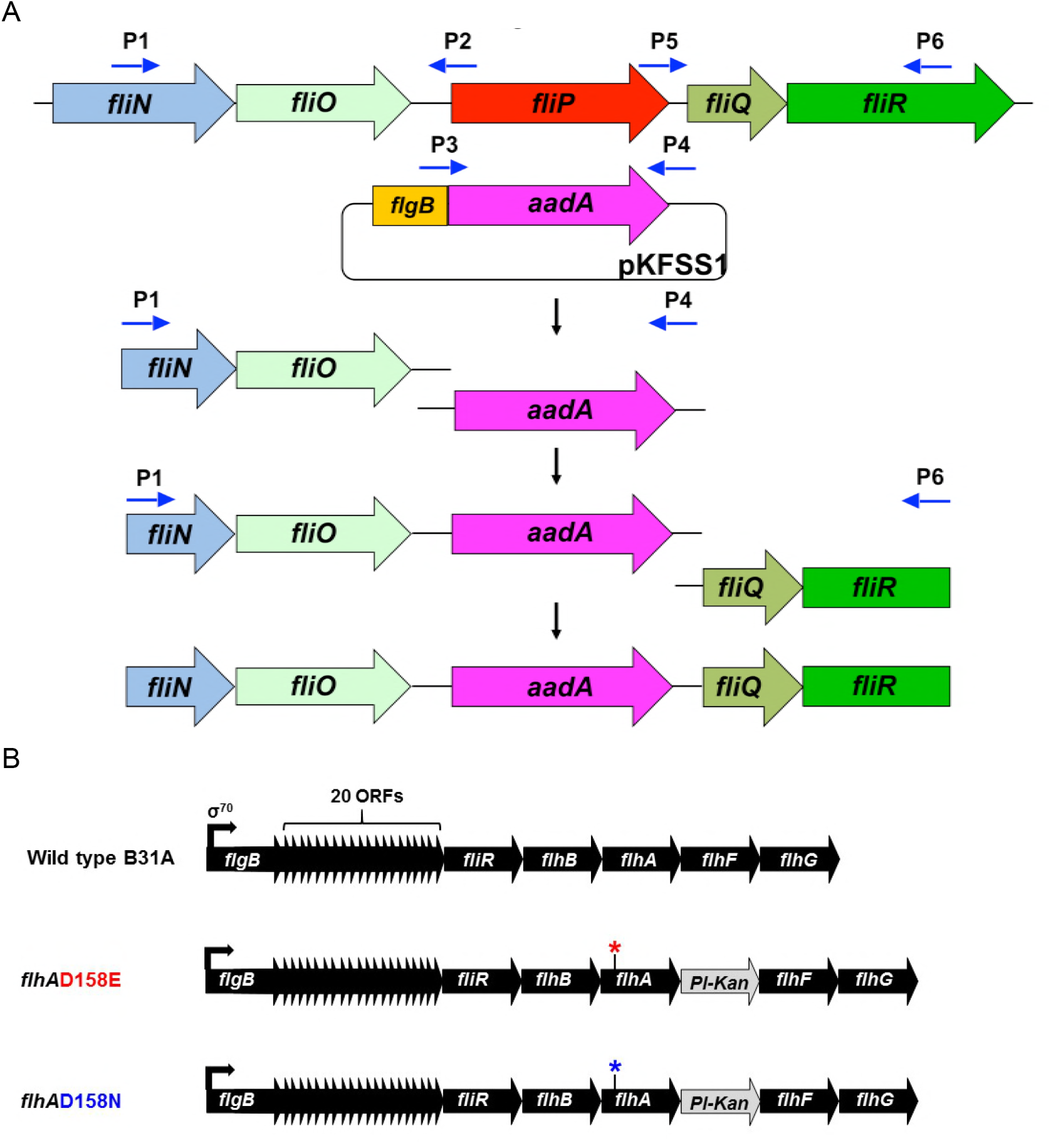
(A) Construction of a non-polar Δ*fliP* mutant in *B. burgdorferi*. Targeted inactivation of *fliP* is achieved by homologous recombination using a streptomycin resistance gene (*aadA*) as a marker. (B) Construction of point mutants in B. *burgdorferi flhA*. Point mutant B. *burgdorferi* clones were created by mutating Asp to Glu or Asn (GAT to GAA or AAT, respectively) at position 158 of FlhA. The point mutant *flhAD158E* exhibits non-motile phenotype whereas the *flhADl58N* cells show reduced motility phenotype. These mutants’ flagellar motors are similar to the *flhA*-deletion mutant Δ*flhA* (not shown).

**Figure Sll.**
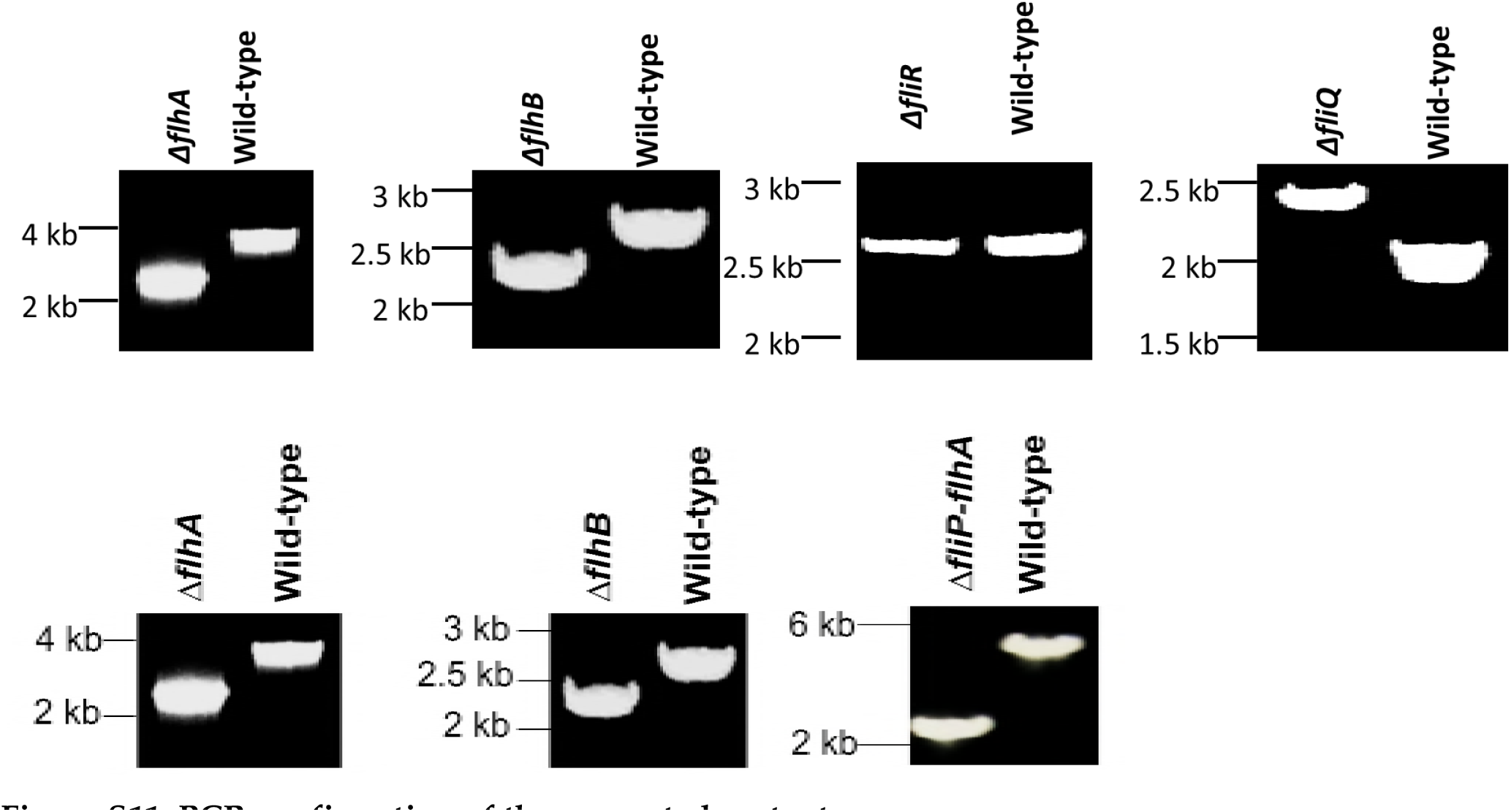
PCR confirmation of the generated mutants. The PCR products are shown as expected.

**Figure S12.**
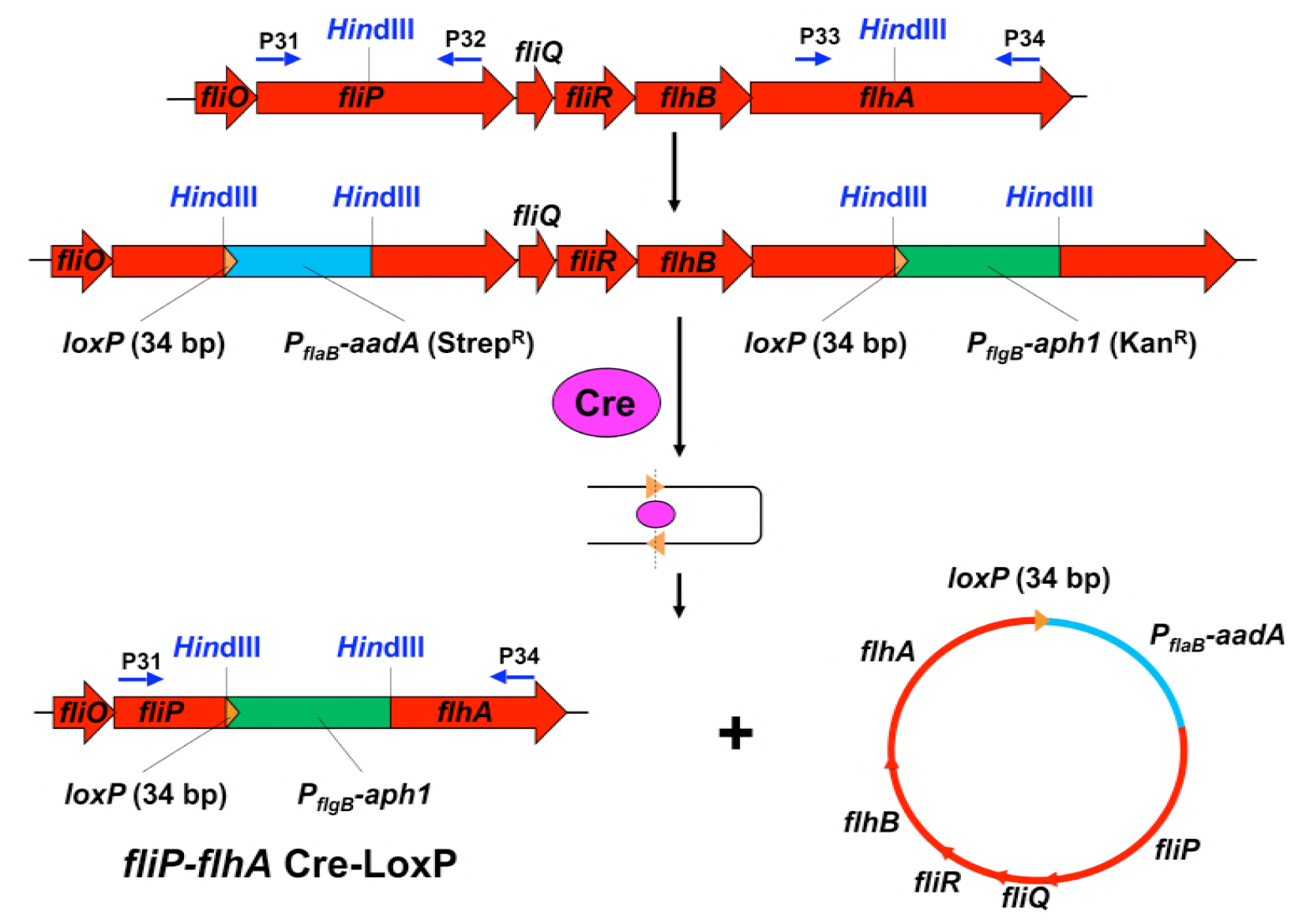
Construction of a quintuple *ΔfiP-flhA* mutant in *B. burgdorferi*. Deletion of the *fliP, fliQ, fliR, flhB, and flhA* genes is achieved by utilizing the Cre-*lox* recombination system.

**Figure S13.**
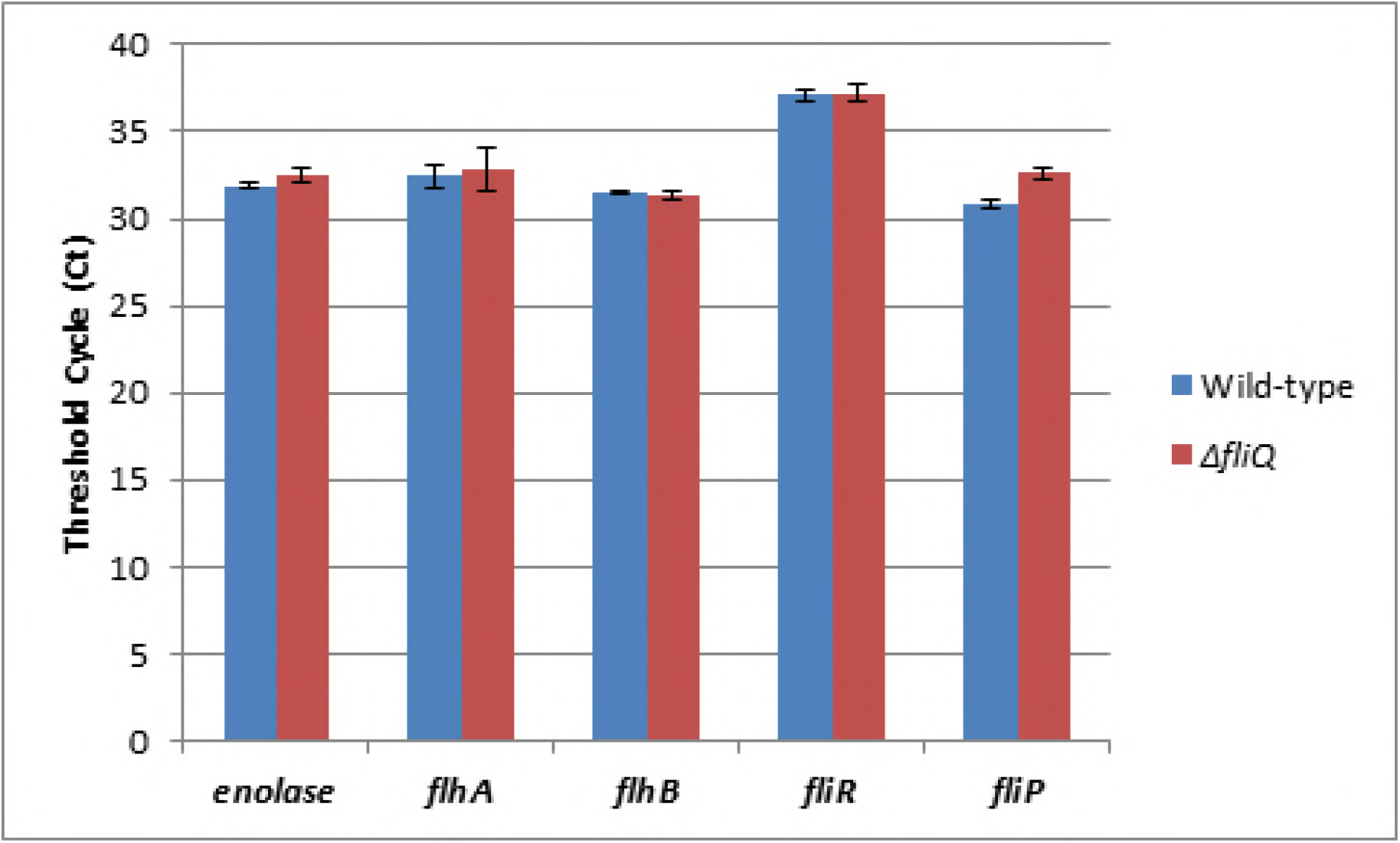
Comparison of transcript levels between wild type and *ΔfliQ*. Transcript levels of *fliP* (which is located downstream of *fliQ*) and *fliR, flhB*, and *flhA* (located upstream of *fliQ*) were measured using RNAs extracted from wild-type and Δ*fliQ* mutant cells. qRT-PCR was performed using those RNA samples. *B. burgdorferi enolase* was used as a control. qRT-PCR were carried out in triplicate and presented as the average of all three data sets (threshold cycle; Ct).

**Figure S14.**
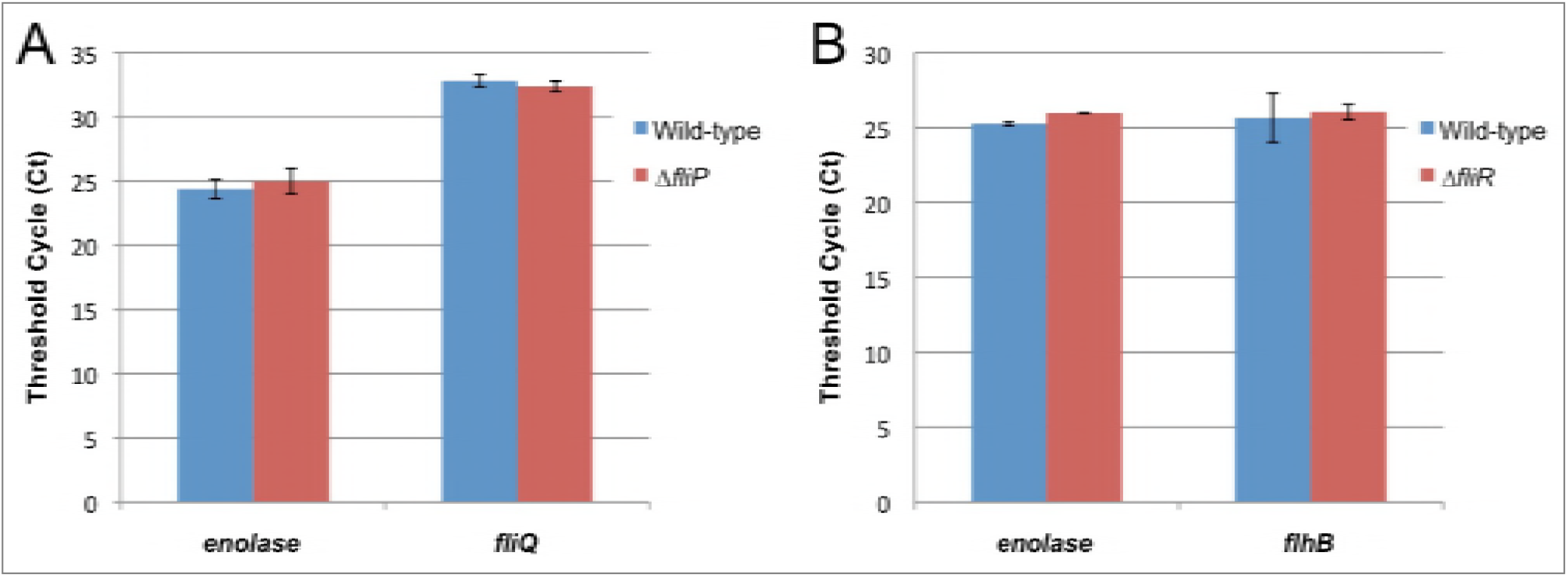
Comparison of transcript levels between wild type, Δ*fliP*, and Δ*fliR*. (A) Transcript level of *fliQ* (which is located downstream of *fliP)* was measured using RNAs extracted from wild-type and Δ*fliP* mutant cells. (B) Transcript level of *flhB* (which is located downstream of *fliR)* was measured using RNAs extracted from wild-type and Δ*fliR* mutant cells. *B. burgdorferi enolase* was used as a control. qRT-PCR were carried out in triplicate and presented as the average of all three data sets (threshold cycle; Ct).

**Table S1.** Sequence comparison of T3S proteins among *B. burgdorferi, S. enterica*, and *B. subtilis*.

| *B. burgdorferi* | *S. enterica* |  | *B. subtilis* |  |
| --- | --- | --- | --- | --- |
|  | IDENTITY | SIMILARITY | IDENTITY | SIMILARITY |
| FlhA | 35% | 60% | 40% | 64% |
| FlhB | 24% | 46% | 32% | 58% |
| FliO (FliZ) | 13% | 28% | 20% | 36% |
| FliP | 41% | 61% | 42% | 64% |
| FliQ | 40% | 66% | 43% | 71% |
| FliR | 19% | 40% | 24% | 45% |
| FliH | 13% | 30% | 20% | 34% |
| FliI | 41% | 63% | 44% | 64% |
| FliJ (FlbA) | 10% | 17% | 13% | 34% |

**Table S2.** Mutants used in this study.

| *B. burgdorferi* | Gene | Gene product | Motility | Reference |
| --- | --- | --- | --- | --- |
| Wild type | N/ A | N/ A | motile | Zhao. *et al.* (2013) |
| Δ*fliO* | BB0276 | Flagellar biosynthesis protein FliO | motile | This study |
| Δ*flhA-flhB-fliP-fliQ-fliR* | N/ A | N/ A | nonmotile | This study |
| Δ*flhA* | BB0271 | Flagellar biosynthesis protein FlhA | nonmotile | This study |
| *flhA* D158E point mutant | BB0271 | Flagellar biosynthesis protein FlhA | nonmotile | This study |
| *flhA* D158N point mutant | BB0271 | Flagellar biosynthesis protein FlhA | less motile | This study |
| Δ*flhB* | BB0272 | Flagellar biosynthesis protein FlhB | nonmotile | This study |
| Δ*fliP* | BB0275 | Flagellar biosynthesis protein FliP | nonmotile | This study |
| Δ*fliQ* | BB0274 | Flagellar biosynthesis protein FliQ | nonmotile | This study |
| Δ*fliR* | BB0273 | Flagellar biosynthesis protein FliR | nonmotile | This study |

**Table S3.**
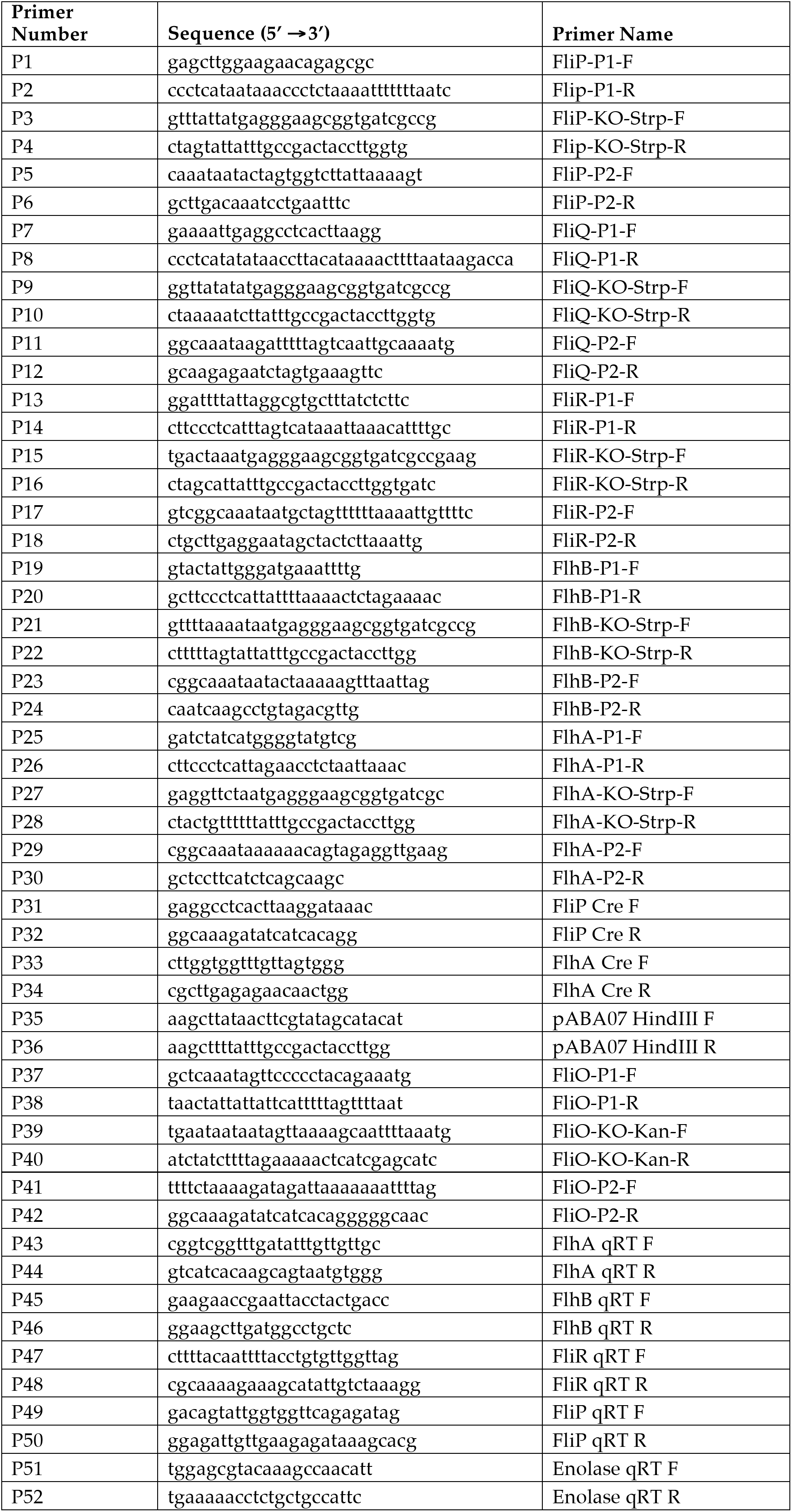
Primers used in this study.

**Table S4.** Cryo-ET and parameters used in this study.

| Genotype | Tomograms | Motors | Magnification | Pixel size (nm) | Resolution (Å) | Accession code |
| --- | --- | --- | --- | --- | --- | --- |
| Wild type | 780 (DDD) | 7,242 | 15,400 (DDD) | 0.25 | 30 | EMD-5627 |
| $\Delta fliO$ | 108 | 546 | 31,000 | 0.57 | 43 | EMD-6090 |
| $\Delta flhA-flhB-fliP-fliQ-fliR$ | 342 | 54 | 31,000 | 0.57 | 63 | EMD-6094 |
| $\Delta flhA$ | 324 | 2,007 | 31,000 | 0.57 | 38 | EMD-6088 |
| $\Delta flhB$ | 543 | 217 | 31,000 | 0.57 | 58 | EMD-6089 |
| $\Delta fliP$ | 265 | 359 | 31,000 | 0.57 | 48 | EMD-6091 |
| $\Delta fliQ$ | 255 | 1,403 | 31,000 | 0.57 | 41 | EMD-6092 |
| $\Delta fliR$ | 229 | 830 | 31,000 | 0.57 | 41 | EMD-6093 |

**Movie S1**. A class average shows the 16-fold symmetry at collar stator region, 23-fold symmetry of the spoke-link links, and 6-fold symmetry of the ATPase complex. The left is the sideview with yellow line slicing through, the right is the cross-slicing view corresponding to the yellow line.

**Movie S2**. Surface rendering of a rebuild map and atomic model fitting. The top part (collar, stator and MS-ring) structure was from global average applied with 16-fold symmetry. The bottom part (C-ring, spoke links and ATPase) structure was from a combine of class averages. Same in Fig. 5 and Fig. 6. In order to refine the bottom structure, the top part of those class averages were not aligned and do not show the 16-fold symmetry. The hexagonal “hub” were segmented in chimera, and the FliI-FliH atomic structure (pdb:5B0O) were initially fitted into the segmented density. However, as there is extra density, and 3 to 4 links extend from the hub, we postulate there are more than one FliH_2_. Three more FliH_2_ were placed next to the FliI-FliH atomic structure, and they were fitted into the density using Molecular Dynamic Flexible Fitting in NAMD (Phillips et al., 2005). The movie here showed the fitting process by morphing. The FliJ (pdb:3AJW) atomic model was inserted to the center of FliI hexamer. The atomic structure of FlhA homolog MxiA (pdb: 4A5P) was fitted into the density by rigid fitting in Chimera. The density of FlihA here was from refined alignment on spoke-like links and thus did not show 9-fold symmetry. The atomic structure for FliG-FliM-FliN was a homology model based on FliGMC-FliMM complex from *Thermotoga maritime* (Vartanian et al., 2012), with the hydrophobic patch of FliN tetramer facing the links. There are 46 copies of FliG-FliM-FliN and they fit reasonably in the C-ring density.

**Movie S3**. The symmetry of spoke-link mismatches with the symmetry of stators, suggesting the C-ring and the ATPase rotate together relatively to the stators. The left are two cross section views from stator (top) and link with the ATPase region (bottom), respectively. The right is the rebuild map orientated correspondently to represent the mismatching. Although the figures were arranged as CW rotation (CCW viewed from bottom), the rotation can be either CW or CCW, as they are WT motors so the rotation orientation cannot be determined solely from cryo-ET image processing.

